# A shared code for perceiving and imagining objects in human ventral temporal cortex

**DOI:** 10.1101/2024.10.05.616828

**Authors:** V. S. Wadia, C. M. Reed, J. M. Chung, L. M. Bateman, A. N. Mamelak, U. Rutishauser, D. Y. Tsao

## Abstract

Mental imagery is a remarkable phenomenon that allows us to remember previous experiences and imagine new ones. Animal studies have yielded rich insight into mechanisms for visual perception, but the neural mechanisms for visual imagery remain poorly understood. Here, we first determined that ∼80% of visually responsive single neurons in human ventral temporal cortex (VTC) use a distributed axis code to represent objects. We then used that code to reconstruct objects and generate maximally effective synthetic stimuli. Finally, we recorded responses from the same neural population while subjects imagined specific objects and found that ∼40% of axis-tuned VTC neurons recapitulated the visual code. Our findings reveal that visual imagery is supported by reactivation of the same neurons involved in perception, providing single neuron evidence for the existence of a generative model in human VTC.

**One Sentence Summary:** Single neurons in human temporal cortex use feature axes to encode objects, and imagery reactivates this code.

---

Mental imagery refers to our brains’ capacity to generate percepts, emotions, and thoughts in the absence of external stimuli. This ability pervades many aspects of the human condition. It allows for the generation of visual art (*1–3*), musical composition (*4–7*), and creative writing (*8*– *10*). It subserves efficient planning (*11*, *12*) and navigation (*13–15*) via the simulation of actions and outcomes. It is also the basis for calling to mind a recent experience, person, place, or object, which is a key aspect of episodic memory (*11*, *16–20*). In a clinical setting, uncontrolled mental imagery can contribute to psychological disorders including anxiety, schizophrenia, and post-traumatic stress disorder (*21*).

Perhaps the most consistent and established finding in mental imagery research is that imagery of a given sense co-opts the neural machinery used for perception, in other words imagery is supported by activation of sensory areas. This has been shown explicitly during auditory (*22–25*), olfactory (*26*, *27*), tactile (*28*), speech (*29*), and even motor imagery (*30–34*), though most extensively in visual imagery (*35–46*); damage in or loss of either dorsal or ventral visual pathways (*47*) often leads to parallel deficits in visual imagery (*48–50*). However, these studies lack the spatial resolution to determine whether imagery reactivates the exact same neural populations that support perception or whether instead imagery is subserved by activation of separate circuitry roughly located in the same cortical regions. Even the strongest previous evidence, consisting of single-unit recordings of face-selective neurons reactivating during free recall in human VTC (*51*), could not address whether imagery reinstated the detailed sensory code for objects, as this code was not mapped.

Here, we attempt to shed light on the single-neuron mechanisms of visual imagery by determining the code for visual objects and then examining whether this code is reactivated during imagery. We focused our investigations on ventral temporal cortex (VTC), a large swath of the human temporal lobe dedicated to representing visual objects (*52*, *53*). We recorded single neurons in human patients implanted with electrodes to localize their focal epilepsy (*54*) as they viewed and subsequently visualized visual objects. First, we found that, as in non-human primates (*55–58*), human VTC neurons showed robust visual responses (*51*, *59–62*), and are well modelled by linearly combining features in deep layers of a deep network trained to perform object classification (*63*). We confirmed two consequences of such a code: each neuron had a linear null space orthogonal to its encoding axis, and each neuron responded maximally to synthetic stimuli generated using its encoding axis. Second, we asked subjects to imagine a diverse subset of objects that they had previously seen, while recording responses of visually-characterized VTC neurons. We found that a subset (∼40%) of axis-tuned neurons reactivated during imagery, and the imagined responses of individual neurons to specific objects were proportional to the projection value of those objects onto the neurons’ preferred axes. Together these findings provide evidence for the implementation of a generative model (*64–66*) in human VTC by neurons that represent both real and imagined stimuli.

## Results

### Neurons show diverse category tuning

We examined how human VTC neurons encode visual objects by recording responses of these neurons to a large set of objects with varying features using a rapid screening task. Patients sequentially viewed a series of 500 images (4 repetitions each, 2000 total trials), drawn from face, text, plant, animal, and object categories (Figure 1A, top; Figure S1A for detailed schematic; Figure S2A for stimulus examples). At random intervals a catch question pertaining to the immediately preceding image would appear on the screen. Images stayed on screen for 250 ms and the inter-trial interval was randomized between 100-150 ms. Patients answered 77% of the catch questions correctly, indicating that they carefully attended to the stimuli (Figure 1B).

**Figure 1.**
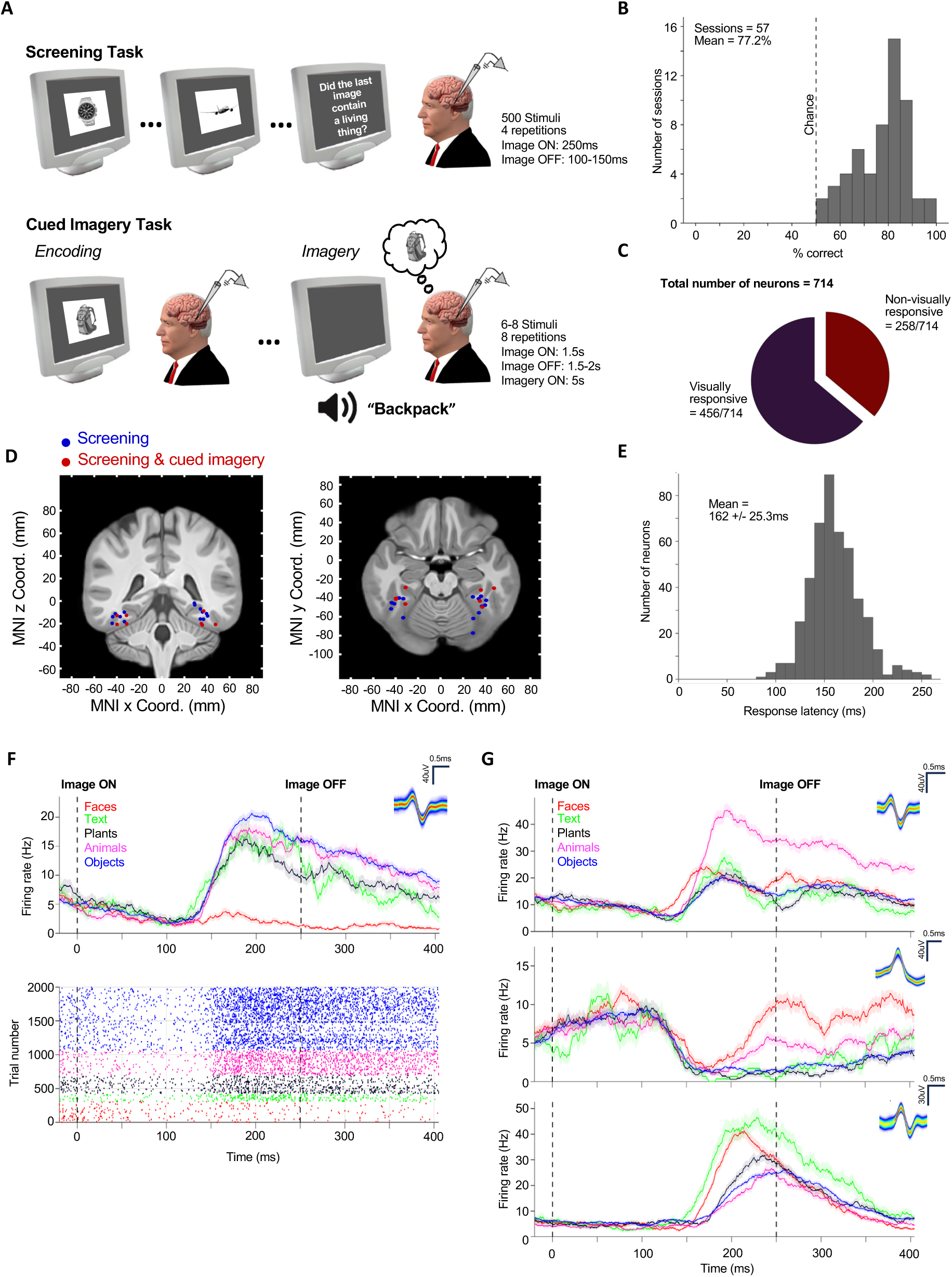
Selective visual responses in human VTC. **(A)** Tasks. Patients performed two tasks: a screening task (top) and a cued imagery task (bottom). In the screening task, 500 grayscale images with white backgrounds were displayed on a gray screen for 250 ms with the inter-trial interval jittered between 100-150 ms. Images subtended 6-7 visual degrees. At random intervals (min interval: 1 trial, max interval: 80 trials) patients answered a yes-no catch question about the image that came before it. In the imagery task, a subset (6–8) of the 500 images used for screening were used, chosen to have spread across both the preferred and principal orthogonal axes. Each trial consisted of 2 images, and an initial encoding period wherein images were displayed on a gray screen for 1.5 s with the inter trial interval jittered between 1.5 – 2 s. After each image was viewed 4 times, a distraction period occurred wherein patients were required to spend 30 s on a visual search puzzle. Finally, after the distraction period they were cued by the experimenter to imagine both images in the trial one by one in alternating order until both had been visualized 4 times. **(B)** Accuracy of catch question responses. Patients answered catch questions correctly in 77 ± 11% of the trials (± SD). Dashed line indicates mean chance level. **(C)** 456 of the 714 recorded neurons were visually responsive. **(D)** Locations of the 27 microwire bundles (left and right) that contained at least one well isolated VTC neuron across all 16 patients. Montreal Neurological Institute coordinates are shown in Table S1. Each dot represents the location of one microwire bundle (8 channels). The locations marked in red were sessions used in the subsequent imagery task. **(E)** Response latencies for the 456 visually responsive neurons relative to stimulus onset (mean 162 ± 25 ms (± SD)). **(F-G)** Example neurons in the screening task. Stimulus onset is at t = 0 and offset is at t = 250 ms. The insets show the mean waveform.

We recorded 714 VTC neurons in 57 sessions across 16 patients. Out of the 714 neurons, 456 showed a response that differed significantly between categories after the onset of a visual stimulus (Figure 1C; 1 x 5 sliding window ANOVA, bin size 50 ms, step size 5 ms, 5 consecutive significant bins with p < 0.01; see methods); we refer to such category tuned neurons as “visually responsive” throughout. The locations of all electrodes from which we recorded can be seen in Figure 1D (red and blue dots); supplementary tables S1 and S2 contain neuron count and patient demographic information. Response onset latency of individual neurons was computed on a trial-by-trial basis using a Poisson burst metric (*67*) (see methods). The mean response latency was 162 ms (Figure 1E). Human VTC neurons showed diverse tuning patterns (Figure 1E-G).

### Neurons are tuned to specific axes in object space

We next examined whether visually responsive neurons encoded specific object features. We leveraged deep networks trained on object classification to build a low-dimensional object space that captures the shape and appearance of arbitrary objects (*68*) without relying on subjective visual feature descriptors. We built our object space by passing the 500 images shown to patients through AlexNet (*63*) and performing principal components analysis (PCA) on the unit activations in layer fc6 (Figure 2A; we chose fc6 because, as in non-human primates (*56*), it explained most neural variance for human VTC neurons, see Figure S4A). 50 dimensions explained 80.68% of the variance in fc6 responses (Figure S2B) and used these 50 dimensions in all analyses unless stated otherwise. This approach allowed us to describe every visual object shown as a point in a 50-dimensional feature space. We next investigated how neural activity mapped onto this feature space.

**Figure 2.**
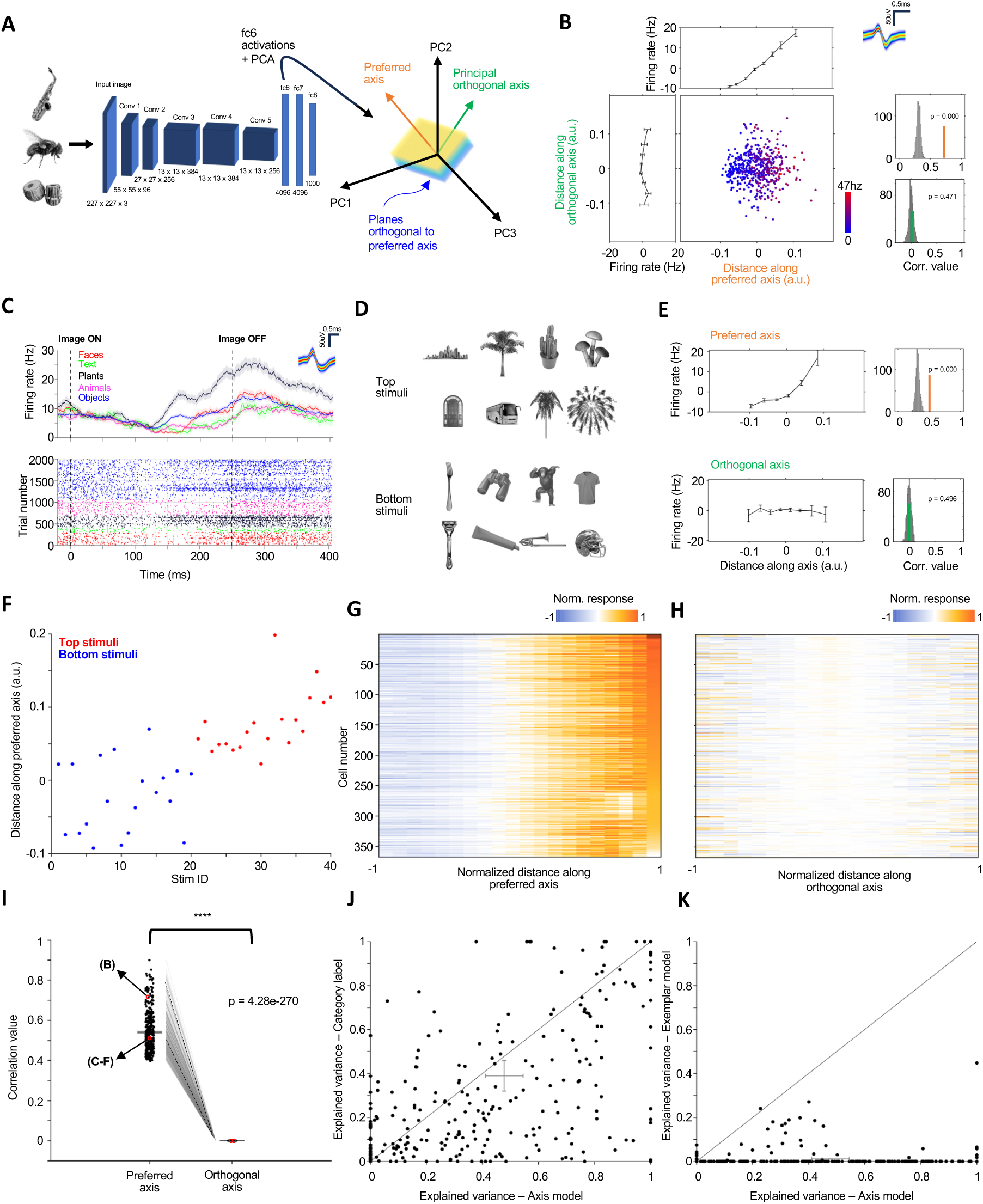
Axis tuning in human VTC neurons. **(A)** Stimulus parametrization and axis computation procedure. Stimuli were parametrized as points in a 50-dimensional feature space formed by the first 50 PCs of the unit activations of AlexNet’s fc6 layer. Neurons’ preferred axes were computed in this space (see methods). **(B)** Example axis-tuned neuron. Scatter shows the mean subtracted neural responses to the 500 stimulus images (color is firing rate) as a function of the projection value of the image on the neuron’s preferred axis and principal (longest) orthogonal axis. (Top) Mean response as a function of distance along the preferred axis. (Left) Mean response as a function of distance along the orthogonal axis. (Top distribution) Mean correlation between projection value onto the preferred axis and firing rate (orange) is significantly larger than expected by chance (p = 0.001, permutation distribution 1000 repetitions). (Bottom distribution) Mean correlation between the projection value onto the orthogonal axis and firing rate (green) is not significantly different from chance (p = 0.496, permutation distribution 1000 repetitions). **(C-F)** Example neuron for which the axis model explains complex neural tuning that does not follow pre-defined categorical boundaries. **(C)** PSTH and raster of the neuron. Raster reveals robust responses to a few out of “preferred” category objects. **(D)** The top (most preferred) and bottom (least preferred) stimuli are not grouped by category. **(E)** Axis tuning. See (B) for notation. **(F)** Projection values of the stimuli shown in (D) reveal a relationship between the projection value and stimulus preference. **(G-J)** Population summary for all axis tuned cells. **(G-H)** Normalized (mean subtracted and scaled by maximum value for each neuron) firing rate as a function of position along the preferred (G) and non-preferred (H) axis. Firing rate increased monotonically along the preferred axis and did not change systematically along the non-preferred axis. Each row is a neuron (n = 367). **(I)** Pearson correlation between the projection value of the stimulus images onto the preferred and orthogonal axes and the firing rate response of each neuron (n = 367). The example neurons are marked in red. **(J)** Model comparison between the axis model and the category model. The axis model explained significantly more variance than the category model (on average 44.92% vs 32.17% explained variance, respectively. p = 5.89e-09, paired t-test). The error bars indicate variance of the mean, each dot is a neuron (n = 318, neurons that had >10% explainable variance, see methods). **(K)** Model comparison between the axis and exemplar model. The axis model explained significantly more variance than the exemplar model (44.92% vs. 1.43% explained variance, respectively. p = 2.44e-67, paired t-test). The error bars indicate variance of the mean.

For each cell, we computed a ‘preferred axis’ given by the coefficients *c*_*pref*_ in the equation 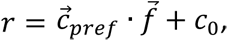 where *r* is the response of the neuron to a given (single) image, 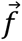 is the 50D object feature vector of that image, and *c*_O_ is a constant offset (see methods). The axis tuning of an example neuron is shown in Figure 2B. This neuron showed a monotonically increasing response to stimuli with higher projection values onto its preferred axis (see methods) while showing no change in tuning along an axis orthogonal to its preferred axis. We confirmed that the correlation between projection value and firing rate was significant when compared to a shufled distribution along the preferred axis (p < 0.01, Figure 2B, top distribution) and not significant along the principal orthogonal axis (p = 0.496, Figure 2B, bottom distribution). This “axis code” emphasizes the geometric picture that VTC neurons encode specific preferred axes in feature space and respond to incoming images, formatted as points in this feature space, in a manner proportional to the image’s projection value onto their preferred axis (*55–58*).

Axis tuning also clarified the response pattern of some neurons with complex tuning. The example neuron in Figure 2C responded most strongly to the plant category, but also responded strongly to specific stimuli in other categories (see raster, Figure 2C). Indeed, no semantic label could obviously delineate between the stimuli that elicited the strongest and weakest response from the neuron (Figure 2D). However, this neuron showed significant axis tuning (Figure 2E), and sorting the stimuli according to their projection value onto the preferred axis shows that the top stimuli were those with the highest projection values and vice versa (Figure 2F). Across the population, we found that a majority (367/456, ∼80%) of visually responsive neurons were significantly axis-tuned (Figure 2G, H), showing a significant positive correlation between projection value and response along the preferred axis with no such correlation along the orthogonal axis (Figure 2I; *r*_*pref*_ = 0.54, *r*_*ortho*_= -1.3833e-10, p = 4.28e-270, paired t-test).

We subsequently compared the variance explained by the axis model to that explained by two alternative models: a category label model and an exemplar model. The category label model was chosen for comparison due to the robust category responses seen in VTC (Figure 1F, G). The exemplar model is a well-known alternative to the axis model (*69–71*) which posits that neurons have maximal responses to specific exemplars (i.e., specific points in object space) and decaying responses to objects with increasing distance from the exemplar (see methods). Examining the 318 neurons with > 10% explainable variance (see methods) showed that only 31/318 neurons had good fits (positive explained variance) across all models (39/318 exemplar model, 282/318 category label, 288/318 axis model) and the axis model explained significantly more variance than either alternative (Fig. 2J, K, percentage of explained out of explainable variance, estimated out of sample, for all models is as follows: 43.39% axis, 38.91% category, p = 5.89e-09, paired t-test; 43.39% axis, 1.14% exemplar, p = 2.24e-67, paired t-test; also see Figure S9 for further comparisons between the axis model and additional category models).

We built our low-dimensional object space by leveraging AlexNet, a deep network trained to perform object classification. To examine whether our finding of axis tuning is dependent on the specific choice of AlexNet we repeated the same analyses using several additional deep neural network models (VGG-16/19/Face, CORNet models, and a vision transformer trained without labels (*72*); see supplementary methods). These analyses revealed that axis tuning in feature spaces built from the fully connected layers of these models similarly explained a large amount of the explainable neural variance (∼45%) in the 367 axis-tuned VTC neurons (Figure S3, S4). Thus the axis model is independent of the specific deep network used to build the feature space.

### Reconstructing objects using human VTC responses

We next investigated the richness of the VTC representation by attempting to reconstruct objects using VTC responses. A consequence of the linear relationship between VTC neuron responses and object features is the ease of learning a linear decoder that predicts object feature values from the population activity (*73–75*). The responses of the neurons can be approximated as a linear combination of object features, with the slopes of the ramps corresponding to the weights (*55*). Then for a population of neurons, 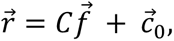 where 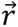 is the response vector of the different neurons, *C* is the weight matrix, 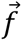 is the vector of object feature values, and 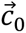 is the offset vector. Inverting this equation to solve for 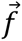 provides a linear decoder that predicts object feature values from the population activity 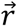 (*73–75*) (see methods). After pooling axis-tuned neurons across all patients (n = 367), we used this approach coupled with leave-one-out cross-validation to learn the linear transform that maps responses to features (Figure 3A).

**Figure 3.**
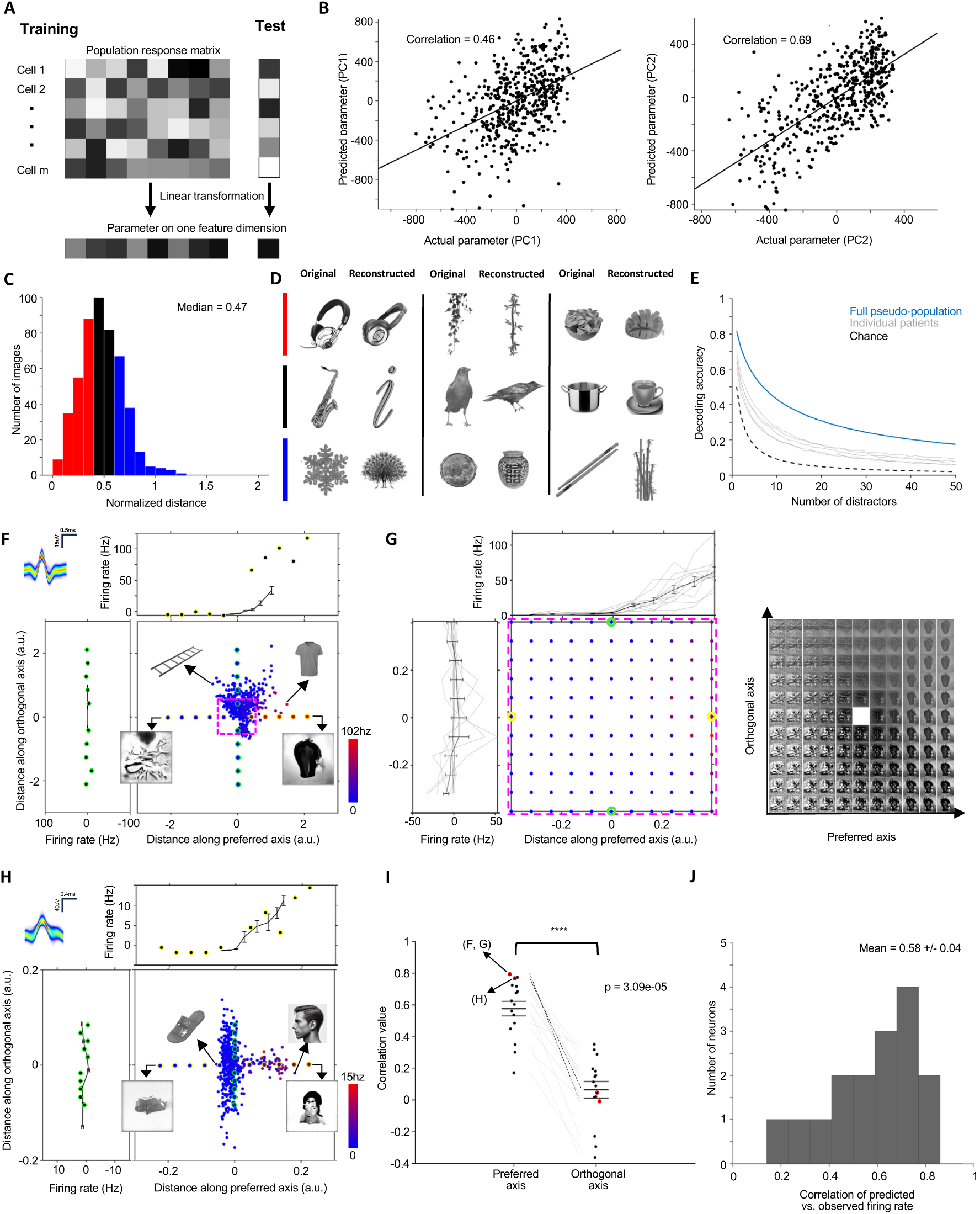
Reconstruction of stimuli from neural activity during viewing and construction of super stimuli. [Note: The human face in panel H of this figure has been replaced with a synthetic face generated by a diffusion model (*130*), in accordance with the bioarxiv policy on displaying human faces.] **(A-E)** Reconstruction of objects from neural responses. **(A)** Decoding model. We used responses to all but one object (500-1 = 499) to determine the transformation between responses and feature values by linear regression and then used that transformation to predict the feature values of the held-out object. **(B)** Decoding accuracy. Predicted vs. actual AlexNet feature values for the first (left, p = 2.6e-28, paired t-test) and second (right, p < 0.001, paired t-test) dimensions of object space (each dot is an image, n = 500, see methods). **(C)** Quantification of decoding accuracy. Normalized distances between reconstructed feature vectors and the feature vectors of the best-possible reconstructions (see methods). A normalized distance of <1 means the decoder found a solution better than chance. The median value is 0.47, and 487/500 images have a distance < 1. **(D)** Example original and reconstructed images, ordered by reconstruction quality. **(E)** Decoding accuracy as a function of the number of distractor objects drawn randomly from the stimulus set (see methods). The black dashed line represents the decoding accuracy expected by chance. The grey lines indicate decoding performance within individual patients. **(F-I)** Construction of to verify the axis model. **(F-G)** Example neuron 1. Yellow and green dots show the positions of the generated super stimuli for which the predicted and observed neural response was compared. The vertical and horizontal line plots show the binned firing rate of the neural response during screening, with the response to the super stimuli shown by yellow and green dots. **(G)** Responses of the same neuron to a grid of generated images sampled densely along the preferred and orthogonal axes. The extent of the grid is indicated by the purple bounding box in (F). (right) The stimuli that comprised the grid. The neuron increased its firing rate only if stimuli varied along the preferred axis. **(H)** Example neuron 2. See (F) for notation. **(I-J)** Population summary. **(I)** Pearson Correlation between the projection value of the generated images and the firing rate response of the neuron for the preferred and orthogonal axes. The two example neurons shown in (F-G) and (H) are marked in red. **(J)** Correlations between the predicted and observed firing rates to the synthetic images.

Figure 3B shows the correlation between actual fc6 feature values and model predictions across all images for the first two dimensions of object space; both correlations were significantly positive (*r*_dim 1_ = 0.46, p = 2.6e-28, paired t-test, *r*_dim 2_ = 0.69, p = 0, paired t-test); Figure S2C shows correlations for all 50 dimensions. To reconstruct objects, we searched a large auxiliary object database for the object with the feature vector closest to that decoded from the neural activity. A normalized distance in feature space between the best possible and actual reconstruction was computed to quantify the decoding accuracy across all stimuli (Figure 3C, see methods). Side-by-side comparisons of the original images with the ‘reconstructions’ chosen from the auxiliary database showed striking visual correspondence (Figure 3D). We also confirmed that, when adding the originally viewed image to a set of distractors the decoder picks the correct object more frequently than expected by chance for regardless of number of distractors (Figure 3E), which is also possible within patient (Figure 3E, grey lines).

### Predicting responses to synthetic stimuli

To further validate the axis model, we attempted to predict the responses of VTC neurons to images not shown to our patients during the initial screening, in a “closed loop” experiment. We used an initial screening session to compute axes for all neurons recorded. We then used a generative adversarial network (GAN) trained to invert the responses of AlexNet (*76*, *77*) to systematically generate images corresponding to evenly spaced points along both the preferred and orthogonal axes. We then returned to the patient room for a second session and rescreened with the synthetic images added to the original stimulus set. We performed this experiment in 8 sessions across 4 patients which yielded 16 axis-tuned neurons.

We predicted that images with projection values greater than the maximum projection value of the original stimulus images should serve as ‘super-stimuli’ driving the neuron to a higher response than any of the original stimulus images (*77*). Figure 3F shows an example neuron tested with the synthetic stimuli. This neuron’s most preferred stimulus was a t-shirt, while the least preferred stimulus was a ladder (Figure 3F); these two stimuli were also the ones that projected most and least strongly onto the cell’s preferred axis. Synthetic stimuli were sampled evenly along both preferred (yellow dots) and orthogonal axes (green dots) such that the extremes had larger projection values than any of the original stimulus images (Figure 3F). Responses to these synthetic stimuli demonstrated the expected increase along the preferred axis and no significant change along the orthogonal axis (Spearman’s rank correlation between projection value and firing rate for preferred axis 0.8303, p = 4e-3 when compared to shufled distribution, orthogonal axis = -0.1037, p = 0.604; Figure 3F). Moreover, the firing rates to the subset of synthetic stimuli with larger projection values along the preferred axis than the original stimulus images were substantially higher than those to the original stimulus images. The most and least effective synthetic stimuli showed striking resemblance to the most and least effective original stimuli. Figure 3G shows responses of the same cell to synthetic stimuli evenly sampled in a 2D grid spanned by the preferred and orthogonal axes (purple square, Figure 3F); responses showed the expected changes in tuning along directions parallel to the preferred axis. Figure 3H shows a second example neuron which was face selective (*r*_*pref*_= 0.89, p = 5.53e-04; *r*_*ortho*_= 0.52, p = 0.12).

We next compared the correlation between firing rate and projection value of the synthetic stimuli along preferred vs. orthogonal axes. Across all 16 neurons the correlation was significantly higher along the preferred axis (Figure 3I, *r*_*pref*_= 0.58, *r*_*ortho*_= 0.08, p = 3.09e-05, paired t-test). We also used the preferred axis to predict the firing rate responses to the synthetic stimuli. The distribution of correlation values between responses predicted using the axes computed from the original stimulus images in the first session and responses to the synthetic images recorded in the second session are shown in Figure 3J (mean = 0.58).

### Neuronal activity during imagery

So far, we have established that human VTC uses an axis code to represent objects and confirmed this code through stimulus reconstruction and generation of super-stimuli. Armed with this understanding of the sensory code, we could now tackle the single neuron mechanisms of visual imagery. In 12 sessions across 6 patients, we tested patients on an imagery task following visual screening (Figure 1A, bottom; Figure S1B for detailed schematic). In the imagery task, patients viewed and subsequently visualized from memory 6-8 objects out of the original 500 used for screening. Each trial required alternating, cued visualization of two object stimuli after an initial encoding period during which patients passively viewed the two stimuli (Figure 1A, bottom, Figure S1B, top). During the imagery period patients would close their eyes and imagine the 2 objects in the trial in alternating 5 s periods until each stimulus had been imagined 4 times, being informed to switch images at the end of a given 5 s period by the experimenter. Each image appeared in 2 separate trials for a total of 8 imagery repetitions per image (see methods). We recorded 231 VTC neurons in the imagery task, of which 131 were visually responsive and 107 were axis-tuned.

We found robust activation of neurons in human VTC during imagery, with 66/231 neurons (∼30%) showing activation to at least one object (Figure S5A, 1 x n sliding window ANOVA or sliding window t-test, n = number of stimuli (6–8), bin size 1.5 s, step size 300 ms, 6 consecutive bins with p < 0.025 for each test – see methods). As in vision, neurons showed a diverse range of response profiles during imagery, some activating sparsely during imagery of a single specific stimulus (Figure S5B), others activating to imagery of multiple stimuli in a graded manner (Figure S5C), and a small number (15/231) activating only during imagery and not during viewing (Figure S5D).

### VTC neurons recapitulate the visual code during imagery

A central goal of the current study was to clarify whether imagery reactivates visual neurons in a way that respects their perceptual code, or whether an alternative code (possibly implemented by a distinct population of neurons) is recruited during imagery. The former would constitute strong evidence for the existence of a generative model (*64–66*) in the human brain. To establish whether VTC neurons reactivate in a manner that respects the axis code, the rapid screening session was conducted using the 500 object images described earlier and axes were computed for the axis-tuned neurons. Then, 6-8 stimuli that were spread along the preferred and orthogonal axes were chosen for use in the imagery task. Figures 4A, B show two example neurons: both showed significant axis tuning and responded most strongly to the image with the largest projection value onto the preferred axis during both viewing (encoding) and imagery (piano and mirror respectively, Figure 4C, D).

**Figure 4.**
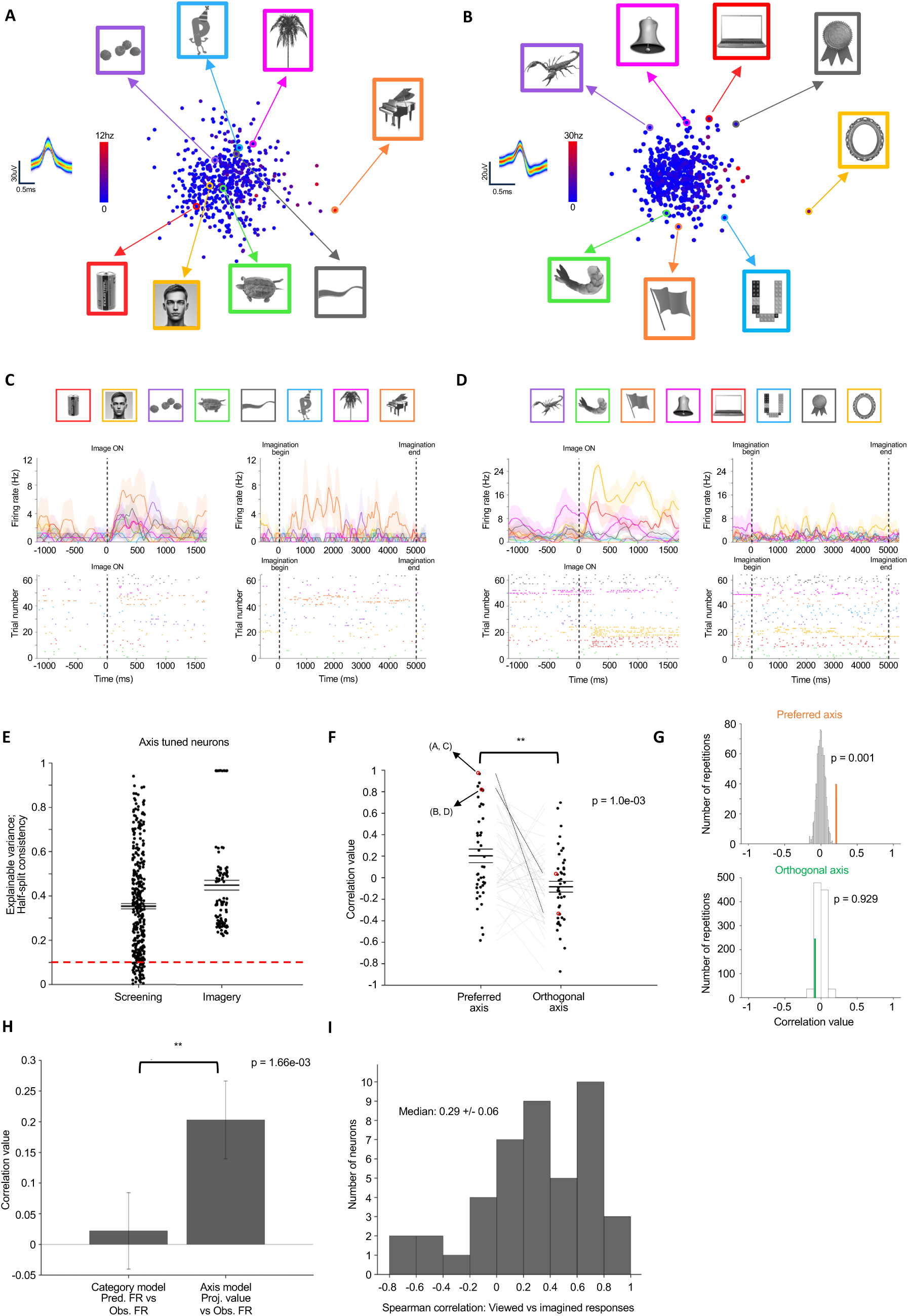
Neurons in VTC reactivate in a manner that respects the visual code. [Note: The human faces in panel A, C of this figure have been replaced with synthetic faces generated by a diffusion model (*130*), in accordance with the bioarxiv policy on displaying human faces.] **(A-D)** Two example neurons. **(A, B)** Axis plots with the subset of 8 stimuli used for imagery indicated. Inset shows the waveform. **(C, D)** Response during encoding/viewing (left) and imagery (right). The top panel shows the stimuli, arranged in order of increasing projection value along the preferred axis. **(E)** Split half consistency (explainable variance) during screening and imagery for all axis-tuned neurons (n = 367 and 107, respectively; each dot is a neuron). Red line indicates cut-off for including a neuron in the explained variance calculation. **(F-G)** Population summary. **(F)** Pearson correlation between the projection value onto the axis computed during screening and firing rate during imagery for the preferred and orthogonal axes in all axis-tuned and reactivated neurons (n = 43). Correlations along the preferred axes were significantly larger than with the orthogonal axis (*r*_*pref*_ = 0.20, *r*_*ortho*_ = -0.084, p = 1.0e-03 paired t-test). The two neurons shown in (A-D) are marked in red. **(G)** Comparison of the mean correlation across all reactivated neurons to the null distribution. Responses were significantly correlated with the projection value onto the preferred axis (p = 0.001, shufled distribution with 1000 repetitions) but not with the projection value onto the orthogonal axis (p = 0.929, shufled distribution with 1000 repetitions). **(H)** Model comparison. The axis model predicted firing rates during imagery significantly better than the category model (error bars are ± SEM., p = 1.66e-3, paired t-test, n = 43) **(I)** Spearman correlation between the mean firing rates during viewing and imagery of each stimulus was positive (0.29 ± 0.06, significantly larger than 0, p = 2.40e-3, one sample t-test, n = 43).

Responses to imagined objects were consistent and reliable, with half-split consistency comparable to that of visual responses during viewing (Figure 4E, mean explainable variance in neurons with explainable variance > 10%; screening = 39.70%, imagery = 44.92%). Examining the population of axis-tuned neurons that reactivated during imagery (43/107, ∼40%) revealed a significant correlation between projection value onto the neurons’ preferred axes (computed using screening responses) and responses during imagery (*r*_*pref*_= 0.20, Figure 4F left, p = 0.001 as compared to a shufled distribution, Figure 4G top) with no significant correlation along the orthogonal axes (*r*_*ortho*_ = -0.08, Figure 4F right, p = 0.929, Figure 4G bottom). When we compared the correlation between projection value and firing rate during imagery for the axis model against the correlation of predicted and observed imagery responses for the category model, we found the former performed significantly better (Figure 4H, p = 1.66e-3, paired t-test; see Table S3 for other models). Lastly, the distribution of Spearman’s rank correlation coefficients between screening and imagery of the same images in reactivated axis-tuned neurons showed a median value significantly larger than 0 (0.29 ± 0.06, p = 2.4e-03, one sample t-test; Figure 4I).

### Decoding of imagined objects

Training a decoder on one task and testing it on another is a powerful way to examine whether two tasks share a common code (cross-condition generalization) (*78*). We next used this approach to further examine the correspondence between the neural code during vision and imagery. We first confirmed that we could decode the identity of imagined objects when training a decoder on imagery responses (Figure 5A). We then trained a decoder on visual responses and tested on imagery responses (Figure 5B). This decoder performed well above chance (all neurons = 31.35%, responsive neurons = 28.12%, chance = 16.67%), confirming the generalization of the code between perception and imagery.

**Figure 5.**
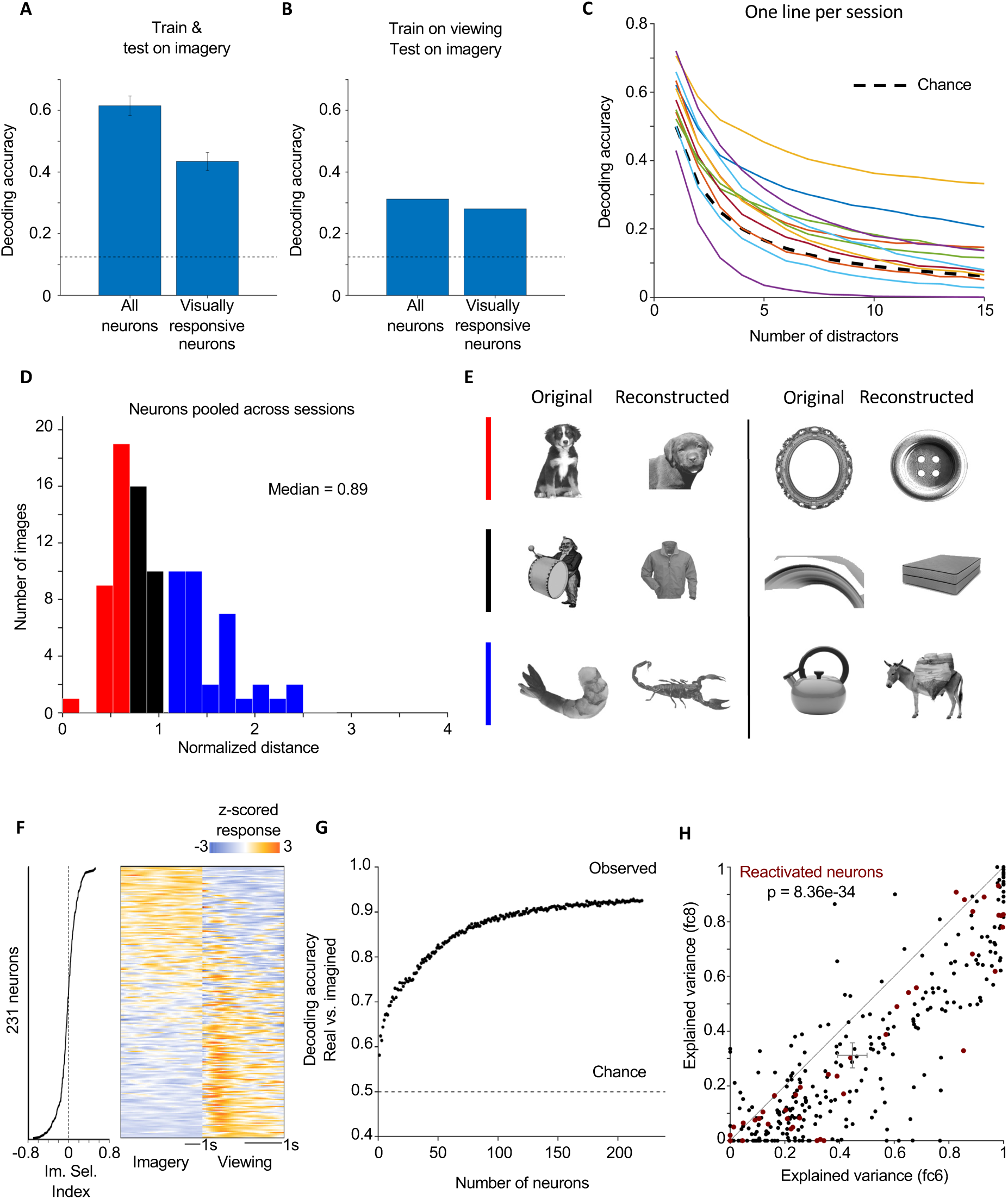
Decoding of imagined object features. **(A)** Image identity could be decoded from neural activity during imagery using all neurons and just visually responsive neurons (p = 4.52e-121, and p = 5.53e-101, one sample t-test vs. chance, respectively). Black line indicates chance. **(B)** Decoders trained on neural activity during viewing and tested on neural activity during imagery were able to decode image identity (all neurons = 31.35%, responsive neurons = 28.12%, chance = 12.5%), showing that the underlying neural code generalized. Black line indicates chance. **(C)** Decoding accuracy as a function of the number of distractor objects drawn randomly from the stimulus set (see methods). Black dashed line is chance, colored lines are different sessions. **(D)** Normalized distances between reconstructed feature vectors and the best-possible reconstructed feature vectors for all imagined images. 53/94 images had a distance < 1. **(E)** Example of reconstructed (right) and imagined (left) images. (**F-G**) Reality monitoring. (**F**) Right: Mean response time courses of all neurons during the encoding and imagery periods of the cued imagery task (n = 231) computed using a sliding window in each condition (see methods) and z-scored. Left: Imagery selectivity index for each neuron. The dotted line marks ImSI = 0 where the neuron responds equally on average in the two task periods. Neurons were ordered by index value. **(G)** Average accuracy of reality decoding (viewing vs. imagery) as a function of number of neurons. Neurons were chosen at random. The decoding performance plateaued at ∼100 neurons. **(H)** Comparison of neural variance explained between visual (fc6) and semantic (fc8) representations in AlexNet. Fc6 explains significantly more variance (48.52% vs. 32.56%, estimated out of sample, p = 8.36e-34 paired t-test). Each dot is a neuron. Neural activity of most neurons (36/43) that were reactivated during imagery was better explained by fc6 than fc8 (maroon dots).

We further examined if neural activity during imagery allows reconstruction of imagined objects. To test this, we learned a linear mapping between the stimulus features and the firing rates during imagery for all but one image and examined how well the visual features of the left-out image were predicted. We did so for all sets of simultaneously recorded, axis-tuned, and reactivated neurons, using the first 5 PCs of fc6 (see methods). For 9/12 sessions, a nearest neighbor search in fc6 for the closest image to that of the fc6 vector predicted by the decoder revealed the correct target image with above chance accuracy regardless of the number of distractors (Figure 5C). Reconstructed images were highly similar to the target images upon visual inspection (Figure 5E), and this was confirmed by quantitative analysis (Figure 5D, median = 0.89). Note that for 64/94 stimuli that were reconstructed, the best possible reconstruction was visually similar but semantically distinct. Taken together, these findings indicate that neurons in human VTC support visual imagery by reinstating the visual axis code used for perception.

If imagery is supported by activity of the same neurons that support perception, a question of interest is: can VTC neurons distinguish between a viewed vs. imagined object (“reality decoding”) (*72*)? If so, how many neurons are needed? Figure 5F shows mean responses across all trials during visual perception and imagery in all neurons recorded in both conditions. Neurons are sorted according to how selective their response is between imagery and viewing (imagery selectivity index, see methods). Some neurons respond more during perception, some more during imagery, but on average the firing rates were not significantly different (1.99 ± 2.07Hz (± SD) in imagery, 2.08 ± 2.15Hz (± SD) in viewing; p = 0.071, two tailed paired t-test, n = 231). Training a decoder to distinguish between visual perception and imagery (see methods) revealed that decoding was possible with high accuracy (Figure 5G), and possible well above chance even for a single randomly chosen VTC neuron (Figure 5G, number of neurons = 1). The decoding accuracy plateaued after ∼100 neurons (Figure 5G). While the range of imagery selectivity index values for all neurons (n = 231) is large [-0.8 0.8] (Figure 5F, left), the range for reactivated axis-tuned neurons that share a code is small and centered on 0 (0.02 ± 0.18, (± SD)), implying that VTC subserves both behaviors (imagery via reactivation and reality monitoring) via separate populations of neurons.

### The axis code is visual in nature

An important question is whether the reactivation we observed was visual or semantic in nature. Many studies implicate VTC in semantic processing (*79–84*) in addition to encoding visual features of objects. To address this question, we first examined the visual vs. semantic content of different layers of AlexNet. Layer fc8 (after SoftMax) harbors the most explicit semantic representation because the 1000 activations in this layer correspond to normalized probabilities of the image belonging to each of the 1000 ImageNet classes (*63*), making the representation categorical and the most semantic by definition. We next asked which layer best explains neural activity of human VTC neurons during viewing and found that fc6 explained significantly more variance (fc6 = 48.52%, fc8 = 32.56%, estimated out of sample, p = 8.36e-34 paired t-test, Figure 5H, S4A). This was also the case when only considering the axis tuned neurons that reactivated during imagery (maroon dots, Figure 5H). Layer fc6 activity also best predicted firing rate during imagery, compared to all other layers (see Table S4). As a further control, we compared the amount of neural variance explained for each neuron by a feature space that is purely visual or purely semantic by construction. For the former, we used a vision transformer (DINO-ViT) trained in an unsupervised manner without labels (*72*), and for the latter, we used a semantic embedding space (GloVe 840B) (*85*) (see supplementary methods and Figure S9). This comparison revealed that the visual model explained significantly more variance than the semantic model (Figure S10B; DINO-ViT = 53.12%, GloVe 840B = 44.23%, p = 9.14e-19 paired t-test). Overall, these analyses demonstrate that the axis code shared by perception and imagery in human VTC is visual and not semantic.

## Discussion

In this paper, we explore the long-standing hypothesis that mental imagery is supported by reactivation of sensory areas in the visual domain. We focused our investigations on VTC, a region long known to harbor representations of complex visual objects (*52*, *53*, *55*, *56*, *58*). We first mapped out the sensory code for visual objects in VTC, and then measured responses of the same population during imagery. Human VTC neurons (367 out of 456 visually responsive neurons recorded) represented visual objects via linear projection of incoming object vectors onto a specific ‘preferred’ axis in a high dimensional feature space built using the unit activations of a deep network. This axis model implies VTC can very efficiently represent all possible objects provided there are enough neurons with independent axes to span object space. Confirming the axis model, we could reconstruct viewed objects from neural activity with high accuracy using a linear decoder (Figure 3A-E) and generate synthetic super-stimuli for cells using the axis mapped to real stimuli (Figure 3F-J).

A subset of neurons (66/231 total, 43/107 axis-tuned) reactivated during imagery in a manner that respected the axis code: imagined responses were significantly correlated with the projection value of the imagined objects onto the preferred axis but not the orthogonal axis (Figure 4G), and viewed and imagined responses were positively correlated (Figure 4I). Our findings demonstrate for the first time that neurons in VTC support visual imagery by reactivating in a manner that respects the visual code. This correspondence between viewing and imagery allowed us to predict neural activity during imagery (Figure 4H), to decode the identity of the imagined object with decoders trained during viewing (Figure 5B), and to reconstruct the images patients imagined (Figure 5D, E).

While perception and imagery activated a shared code, decoders with access to a random subset of VTC neurons could nevertheless easily distinguish between whether an image was viewed or imagined (Figure 5G). Thus, processes downstream of VTC can do the same (*40*, *42*, *78*). All patients tested scored highly in the vividness of visual imagery questionnaire, so this study cannot comment on how these results would differ in subjects with dim or absent visual imagery.

We determined that the axis code being used by VTC neurons and shared between viewing and imagery is predominantly visual, because (1) across different layers of AlexNet, neural responses were best explained by visual feature-dominated fc6 and (2) across different deep network models, neural responses were best explained by DINO-ViT, which is purely visual by construction (Figure S10B) (*86–93*).

The source of the top-down signal driving VTC reactivation during imagery remains an open question. Candidates for this source include the hippocampus and prefrontal cortex, given their involvement in various forms of memory (*94*, *95*), their dense connections to VTC (*19*, *96*, *97*), and the known ability of human hippocampal neurons to be selectively reactivated by free recall (*19*, *96*, *97*). Another question is the relationship between the VTC signals during imagery and those previously reported in primary visual areas (V1/V2) (*94*). Given hierarchically organized feedback connections (*65*, *100*, *101*), we hypothesize that the VTC signals may be driving the imagery-related signals in earlier visual areas. Future work is required to investigate the response characteristics of reactivated neurons across the brain, including those both upstream and downstream of VTC during imagery.

This work is the first to reveal a detailed understanding of the neural codes underlying visual object perception and imagery in human VTC. In particular, the demonstration that imagery and perception share a common visual code provides evidence for the existence of a generative model in the human brain—a mechanism by which neurons in the visual pathway reactivate during imagery to enable the synthesis of detailed sensory contents (*65*, *100*, *101*). Generative models solve core challenges for intelligent systems, such as unsupervised learning (*64*) and inference under ambiguity (*102*). The existence of a generative model in the human brain may even explain creative artistic processes that have so far remained out of reach of neuroscientific understanding.

## Methods

### Participants

The study participants were 16 adult patients who were implanted with depth electrodes for seizure monitoring as part of an evaluation for treatment of drug-resistant epilepsy (see Table S2 for demographic data). All patients provided informed consent and volunteered to participate in this study. Research protocols were approved by the institutional review board of Cedars-Sinai Medical Center (Study 572). The tasks were conducted while patients stayed in the epilepsy monitoring unit following implantation of depth electrodes. The location of the implanted electrodes was solely determined by clinical needs. The neural results were analyzed across all 16 patients. Each of the 16 patients included in this study had at least one depth electrode targeting the ventral temporal cortex.

### Psychophysical tasks

Patients participated in three different tasks: an initial screening, cued imagery, and a final re-screening. The initial screening session was conducted to identify axis-tuned neurons. Then 6-8 stimuli were chosen for the imagery task that had some spread along both the preferred and orthogonal axes. After identifying axis-tuned neurons and choosing stimuli, we then conducted the cued imagery task, followed by another screening session immediately after. The second screening was used to match the neurons from the first and second sessions.

#### Screening

Patients viewed 500 object stimuli (grayscale, white background, size: 224×224) with varying features (taken from www.freepngs.com) 4 times each for a total of 2000 trials in a shufled order. Images were displayed on laptop computer with a 15.5” screen placed 1 meter away and subtended 6-7 visual degrees. Each image stayed on screen for 250 ms, followed by an inter-trial interval consisting of a blank screen shown for 100-150 ms (randomized). The task was punctuated with yes/no ‘catch’ questions pertaining to the image that came right before the question requiring the patients to pay close attention in order to answer them correctly (Figure S1A). Catch questions occurred randomly between 2 to 80 trials after the previous one. Patients responded to the questions using an RB-844 response pad (https://cedrus.com/rb_series/). During the synthetic image screens, the synthetic images were added to the original stimulus set.

#### Cued Imagery

Patients viewed a set of 6-8 object stimuli chosen from the 500 used for screening (taken from www.freepngs.com). Each trial focused on 2 images and consisted of an encoding period, a visual search distraction period, and a cued imagery period. During encoding patients viewed the 2 images 4 times each in random order. Each image stayed on screen for 1.5 s and the inter-image interval was 1.5-2 s. After the encoding period a visual search puzzle was presented (puzzles created by artist Gergely Dudas https://thedudolf.blogspot.com/) for 30 s. After reporting via button press whether or not they were able to find the object in the puzzle, patients began the cued imagery period. During cued imagery patients closed their eyes and imagine the stimuli in the trial in an alternating fashion for 4 repetitions of 5 s each (40 s continuous imagery period). Patients were cued to switch the image they were imagining every 5 s by verbal cue (Figure S1B). After the imagery period patients would begin the next trial via button press when ready. Every image was present in 2 trials, leading to 8 repetitions of both encoding and imagery for each image.

##### Workflow

A typical experiment day unfolds as follows: we perform an initial screening session, followed by spike sorting and analysis to assess whether there are visually responsive and axis-tuned neurons in VTC (see below for methods). If there are, we return to the patient room (typically 2-3 hours after the end of the initial screening session) and conduct the cued imagery task immediately followed by the screening task. As a result, each day consists of a cued imagery task, in between two screening sessions. The tasks in the second session (cued imagery and screening) are sorted together so we can assess vision and imagery correspondence in the same neurons. The neurons in the initial and post-imagery screens are matched to remove duplicates (see methods section ‘Matching neurons across experiment sessions’).

### Electrophysiology

The data in this paper was recorded from left and/or right VTC using Behnke-Fried micro-macro electrodes (Ad-Tech Medical Instrument Corporation) (*54*). All analyses in this paper are based on the signals recorded from the 8 microwires protruding from the end of each VTC electrode. Recordings were performed with an FDA-approved electrophysiology system, and sampled at 32khz (ATLAS, Neuralynx Inc.) (*103*).

#### Spike sorting and quality metrics

Signals were bandpass filtered ofline in the range of 300-3000hz with a zero phase lag filter before spike detection. Spike detection and sorting was carried out via the semiautomated template matching algorithm Osort (*104*). The properties of clusters identified as putative neurons and subsequently used for analysis were documented using a suite of spike quality metrics (Figure S6).

#### Electrode localization

Electrode localization was based on postoperative computed tomography (CT) scans. We co-registered postoperative CT onto the preoperative MRIs using Freesurfer’s mri_robust_register (*105*). To summarize recording locations across participants we aligned each participant’s preoperative MRI to the CITI168 template brain in MNI152 coordinates (*106*) using a concatenation of an affine transformation and symmetric image normalization (SyN) diffeomorphic transform (*107*). The MNI coordinates of the microwires from a given electrode shank were marked as one location. MNI coordinates of microwires with putative neurons detected from all participants were plotted on a template brain for visualization (Figure 1D).

### Data Analyses

#### Visual responsiveness classification

To assess whether a neuron was visually responsive we used a 1 x 5 sliding window ANOVA with the factor visual category (face, plant, animal, text, object). We counted spikes in an 80-400 ms period relative to stimulus onset, using a bin size of 50 ms and a step size of 5 ms. Beginning at each time point, the average response in each 50 ms trial snippet was computed and the vector of responses, labeled by their stimulus identity, were fed into the ANOVA. The time point was then incremented by 5 ms and the ANOVA was re-computed. A neuron was considered visually responsive if the ANOVA was significant (p < 0.01) for 6 consecutive time points. These parameters were chosen to ensure that the probability of selecting a neuron by chance was less than 0.05 (compared to permutation distribution with 1000 repetitions, Figure S7A).

#### Response latency computation

We computed single trial onset latencies by using a Poisson spike-train analysis for all visually responsive neurons. This method detects points of time in which the observed inter-spike intervals (ISI) deviate significantly from that assumed by a constant-rate Poisson process. This is done by maximizing a Poisson surprise index (*67*). The mean firing rate of the neuron during the inter-trial interval was used to set the baseline rate for the Poisson process. Spikes from a window of 80-300 ms after stimulus onset were included. For a given burst of spikes, if the probability that said burst was produced by a constant-rate Poisson process - where the rate parameters are specified by the baseline firing rate - was less than 0.001, we took the timepoint of the first spike as the onset latency. The response latency of the neuron was taken to be the average latency across all trials.

#### Response vector computation for individual neurons

All analyses are based on the response of a given neuron to a given image. In the screening task, the response of a neuron to an individual image was computed by averaging spikes in a 250 ms long window that begins at the time of the neuron’s response latency. The cued imagery task consisted of the two periods encoding and imagery (Figure S1B). Encoding responses were computed using a 1.5 s long window beginning at the neuron’s response latency (computed during screening). Imagery responses for reactivated neurons were computed in a 3 s long window beginning at the first time bin the sliding window ANOVA or sliding window t-test was significant (see ‘Reactivation metric’ below). For analyses that used all recorded neurons in the cued imagery task (Figure 5A, B, F, G), encoding spikes were averaged in a 2 s long window beginning at image onset, and across the whole 5 s period for imagery beginning at imagery onset.

#### Object space

We built a high dimensional object space by feeding our 500 stimulus images into the pre-trained MATLAB implementation of AlexNet (*76*) (Deep Learning Toolbox, command: ‘net = alexnet’). The responses of the 4096 nodes in fc6 were extracted to form a 500 x 4096 matrix (using the ‘activations’ function). PCA was then performed on this matrix yielding 499 PCs, each of length 4096. To reduce the dimensionality of this space we retained only the first 50 PCs, which together captured 80.68% of the response variance across fc6 units (Figure S2B). The first two dimensions accounted for 20.17% of the response variance across fc6 units.

#### Axis computation

##### Preferred axis

The preferred axis of each neuron was computed using linear regression. The neural response vector was computed by binning spikes elicited by each stimulus in either a 250ms window starting from the response latency of the neuron (all analyses except model comparisons), or a fixed window 100 ms to 350 ms after stimulus onset (for model comparisons, see ‘Explained variance computation’ section below).

Once the neural response vector was computed the preferred axis is computed as:

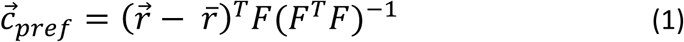

where 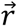 is the *n* x 1 neural response vector to *n* objects, 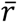 is the mean firing rate, and *F* is an *n* x *d* matrix of features with each row corresponding to the features for a given object that is computed via PCA on deep network activations (see above). The projection value of the stimulus objects onto the preferred axis is given by:

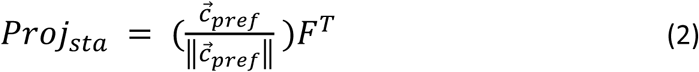

##### Principal orthogonal axis

The orthogonal axis seen in all plots is the principal orthogonal axis. This is defined as the axis orthogonal to the preferred axis along which there is the most variation. This is computed for each neuron as follows: the component along the preferred axis 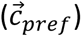 was subtracted from each object feature vector (row) in *F* leaving a matrix of residual feature vectors *F*_⊥_. Succinctly, for a given feature vector 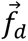 in feature space we computed:

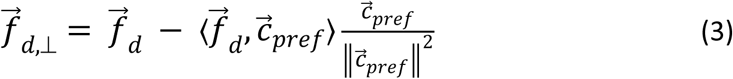

Where 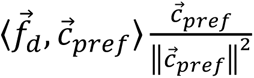is the component of 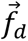 in the direction of 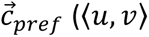 denotes the dot product of vectors u and v). The set of all *n* such vectors (*n* = number of objects) produced an *n x d* matrix *F*_⊥_. Principal component analysis was performed on this matrix *F*_⊥_, and the first principal component was chosen as the principal orthogonal axis.

##### Quantifying significance of axis tuning

For each neuron after the preferred axis was computed we examined the correlation between the firing rate response to the stimuli and their projection value along the preferred axis. This correlation value was recomputed after shufling the features (1000 repetitions) and the original value was compared to this permutation distribution. If the original value was greater than 99% of the shufled values the neuron was considered axis-tuned.

#### Explained variance computation

##### Axis model

The axis model assumes a linear relationship between the projection value of an incoming object onto the neurons preferred axis and its response. Therefore, to quantify the explained variance for each neuron, we fit a linear regression model between the PCs of the features and the responses of the neuron. A leave-one-out cross validation approach was used i.e. the responses to 499 objects were used to fit the model, and the responses of the neuron to the left-out object was predicted using the same linear transform. In this manner we could produce a predicted response for all images. Note that the computation of variance explained by the category label was done in this manner as well, replacing the PC features with the vector of category labels.

##### Exemplar model

The exemplar model assumes that each neuron has a maximal response to a specific exemplar in object space and that the response of the neuron to an incoming object decays as a function of the distance from the object to this exemplar. We used a previous implementation of the exemplar model (*55*) in which the response of the neuron which has an exemplar 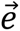 to an incoming object 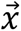 is:

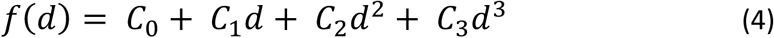

where *d* is the Euclidean distance between the exemplar object and the incoming object:

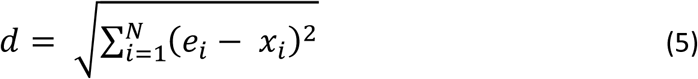

and *N* is the dimensionality of the object space, which in our analyses was 50. In such an implementation the coefficients of the polynomial *C* and the features of the exemplar 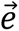 are considered free parameters. They were adjusted iteratively to minimize the error of fit using the MATLAB function *lsqcurvefit*.

##### Category model

The category model posits that the category label of an image is a good predictor of a neuron’s response to that image. To fit this model, we fit a generalized linear model (GLM) with a Poisson link function. The category labels were converted into a one-hot encoded matrix via the MATLAB function *dummyvar,* creating binary indicator variables for each of the five categories. The Poisson GLM was specified as:

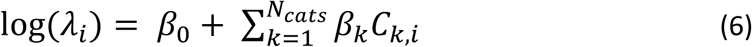

where *λ*_*i*_is the expected firing rate for image *i*, *β*_O_ is the intercept, *β*_*k*_are the category-specific coefficients, *C*_*k*,*i*_ are the binary indicators for category membership, and *N*_*cats*_is the total number of categories considered (equal to 5 except for analysis shown in Figure S9). We used a leave-one-out cross validation approach to evaluate the amount of neural variance explained by this model. To do so, for each neuron, a GLM was fit between responses to all but one image and the binary categorical indicators using the MATLAB function *glmfit.* The predicted firing rate for the held-out image was then computed using the fit *β* values and the held-out category indicator via the function *glmval*.

#### Explainable variance computation

To set an upper bound for the explainable variance, different trials of responses to the same stimuli were randomly split into two halves. The Pearson correlation (*r*) between the average responses from the two half-splits across images was then calculated (averaged across 1000 splits) and corrected using the Spearman-Brown correction:

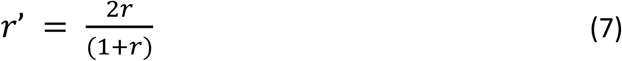

The square of *r*′ was considered the upper bound, which we refer to as the explainable variance of a neuron. The reported results are the ratio of the explained to the explainable variance.

To compare explained variance between different models (Figure 2J, K, Figure S9D-G), all neurons were included without pre-selection. To do so, we computed the response for each neuron in a bin 100 ms to 350 ms after stimulus onset, so that each model had access to exactly the same neuronal response. In addition to variance explained, we also evaluated the proportion of neurons that explained significant amounts of variance (positive explained variance) for each model (Figure S9E, G).

#### Decoding analysis: Screening

Since neurons in human VTC are performing linear projection onto specific preferred axis in object space, their responses can be well modeled as a population by the equation:

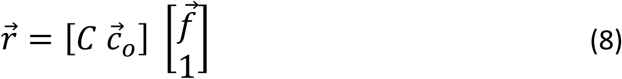

where 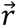 is the *n* x 1 population response vector to a given image (*n* = number of neurons), *C* is the *n* x *d* weight matrix containing the preferred axis for each of the neurons (*d* = number of dimensions, i.e. 50), 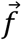 is the *d* x 1 vector of object feature values, and 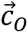 is the *n* x 1 offset vector. Decoding analysis was performed by inverting (8), yielding:

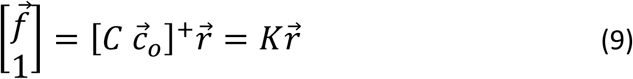

Where *C*^+^indicates the Moore-Penrose pseudoinverse, and *K* is the (*d* + 1) x *n* matrix that transforms measured firing rates 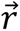 into predicted features 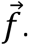 We used the responses of all but one of the objects (500 – 1 = 499) to determine *K* using the MATLAB function *regress*. These were then substituted into (9) to predict the feature vector of the last object. For the *i*^th^ image

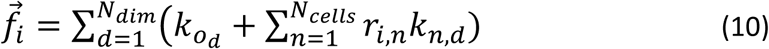

Where *r*_*i*,*n*_is the *n*^th^ element of 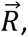 i.e. the response of neuron *n* to the image *i*, *k*_*n*,*d*_is the (*n*, *d*) element of the weight (regression coefficient) matrix, and 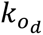 is the *d*^th^ element of the offset vector. Decoding accuracy was quantified by randomly selecting a subset of object images that included the actual feature vector of the decoded object from the total set of 500 and compared their feature vectors to the predicted feature vector of the decoded object by Euclidean distance. If the actual feature vector closest to the predicted feature vector is of the object being decoded (‘target’) the decoding is considered correct. This procedure is repeated 1000 times for each of the 500 images with a varying number of distractors to get an aggregate measure of decoding accuracy (Figure 3E).

#### Object ‘reconstruction’

To generate images that reflect the features encoded in the neural responses we gathered images from an auxiliary database and passed 15,901 background free images through AlexNet. The images were then projected into the space built by the 500 stimulus objects. None of these ∼16k images had been shown to the patients. For each stimulus image the feature vector decoded from the neural activity was compared to the feature vectors of the large stimulus set. The object in the large image set with the smallest Euclidean distance to the decoded feature vector was considered the ‘reconstruction’ of that stimulus image (*56*).

To account for the fact that the large object set did not contain any images shown to the patients, which sets a limit on how good the reconstruction can be, we computed a ‘normalized distance’ to quantify the reconstruction accuracy for each object. We defined the normalized reconstruction distance for an image as

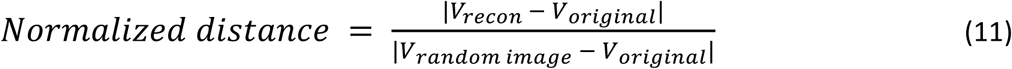

where *V*_*recon*_ is the feature vector reconstructed from neuronal responses, *V*_*original*_is the feature vector of the image presented to the patients, and *V*_*random image*_ is the feature vector of an image drawn at random from the large image set. A normalized distance below 1 means the decoder is performing better than picking an image at random. The median normalized distance value for our data was 0.47. The normalized distance was less than 1 for most (487/500, ∼97%) of the images (Figure 3C), with the distance of the worst performing image being 1.51, implying that the neural responses captured many of the fine feature details of the original objects.

#### Generation of synthetic stimuli

The axis model provides a very clear relationship between images and responses for individual neurons. In essence, images with increasing projection values onto a neuron’s preferred axis will show increasing firing rates. This implies that if one computes a neuron’s preferred axis and then evenly samples points along it and generates images from those points, those images will elicit systematically increasing responses from the neuron. This also implies that if one generates an image from a point further along the axis than any of the stimulus images used to compute the neurons axis, that image will act as a super stimulus and drive the neuron to a higher firing rate than any of the stimulus images.

To test these predictions we ran the screening task in one session, computed the axes for the neurons recorded, sampled points along the preferred and orthogonal axes and fed those vectors back into a pre-trained GAN (*104*) to generate the synthetic stimuli. We then went back to the patient room and re-ran the screening task with the synthetic images added in. The neurons from the first and second sessions were matched (see below) and the responses of the neurons to the synthetic stimuli were recorded.

#### Computation of predicted responses to synthetic images

Predicted responses to the synthetic images were computed by fitting a linear regression model between the PCs of the features and the responses of the neuron during the first session to the 500 stimulus images. That linear transform was then used to predict the responses to the synthetic images. These predicted responses were then compared to the responses recorded in matching neuron during the second session.

#### Matching neurons across experiment sessions

We assessed whether neurons in the initial and post-imagery session were the same using the following procedure. We first compute the selectivity vector (rank ordered list of stimulus number for the neuron), the waveform, the burst index (which is a measure of how many bursts per unit time the neuron discharges (*104*)), the computed response latency of the neuron, and whether or not the neuron was axis-tuned (binary variable). We then compared these ‘selectivity vectors’ using cosine distance (MATLAB function *pdist*) and the waveforms by Euclidean distance as following. Each axis-tuned neuron in the initial screening session was compared to all ipsilateral axis-tuned neurons in the subsequent session (same procedure for non-axis-tuned neurons). The session two neurons were then rank ordered in every metric with first place in a given metric giving a session two neuron a score of *x* where *x* = number of session two neurons being compared to the one session one neuron, second place receiving a score of *x* − 1 all the way until the last place neuron in a given metric receives a score of 1. The scores of all session two neurons were then summed and the algorithm would assign the session two neuron with the max score to be the session one neuron’s ‘match’ (Figure S8A-B illustrates the procedure and Figure S8E-H shows examples). This procedure is then repeated in the reverse direction, i.e. each session two neuron is compared to all session one neurons. Pairs that were bijective were automatically returned as ‘matches’ and all others were marked out for manual curation. Manual curation was carried out by examining the used metrics in addition to shape of the peri-stimulus time histogram of the category response of all potential neuron match pairs. We evaluated the quality of the final matching result by assessing the angles between the axes of paired cells (Figure S8C) and by correlating the mean response to each category between pairs of cells (Figure S8D).

#### Reactivation metric

To assess whether a neuron was active during imagery we used a combination of a 1 x N (N = number of images) sliding window ANOVA and a sliding window t-test relative to baseline (500 ms before imagery onset) during the cued imagery period. We counted spikes in a 0-5 s window relative to imagery onset (i.e. the entire cued imagery period), using a bin size of 1.5 s and a step size of 300 ms. Beginning at each time point, the trial average was computed using spikes in a 1.5 s window and the vector of trials was fed into the ANOVA or t-test. The two tests were used in conjunction to capture the different reactivation types found (Figure S5B-D). Some neurons reactivated ‘sparsely’ to only one or two of the stimuli imagined (Figure S5B, Figure 4C, D), while others were more ‘densely’ reactivated increasing their firing rate response for most or all stimuli imagined (Figure S5C, D). The ANOVA was effective at capturing the former while the t-test was more effective at identifying densely reactivated neurons. For the ANOVA the trials were labeled by their stimulus identity. The time point was incremented by 300 ms and the ANOVA was re-computed. A neuron was considered active during imagery if either the ANOVA or the t-test was significant (p < 0.025 for each test) for 6 consecutive bins (or 5 consecutive steps). These parameters were chosen to ensure that the probability of selecting a neuron by chance (p_ANOVA_ + p_t-test_) was less than 0.05 (compared to permutation distribution with 1000 repetitions, Figure S7B, C).

#### Correlation of viewed and imagined responses

For neurons that were active during imagery we computed the correlation between viewed and imagined responses. To compute the viewed response to each stimulus, we collected spikes in a 1 s window of the encoding period starting from the response latency of the neuron (computed during screening) and averaged across repetitions of a given stimulus. To compute the imagined response for all neurons active during imagery, we collected spikes in a 3 s window starting from the first significant time bin (1 x N sliding window ANOVA or sliding window t-test, N = number of stimuli, p < 0.025 for each test) and averaged across repetitions of a given stimulus. The Spearman rank correlation (*r*) was then computed between these 2 vectors.

#### Decoding analyses: Imagery

##### Decoding stimulus identity: Cross condition Generalization

To assess the stability of neural representations between the different task phases (viewing and imagery), Support Vector Machines (SVM) classifiers with uniform class priors and linear kernels were trained on neural activity during the encoding period of the cued imagery task and then tested on the cued imagery period. Stimulus labels were matched between training and testing conditions to ensure valid comparisons. Training data consisted of spikes binned in a 2000 ms window starting from stimulus onset during encoding and the whole 5000 ms window during imagery. This analysis was done on a pseudo-population of all recorded neurons across imagery sessions.

##### Decoding stimulus identity: Within-condition

To determine whether stimulus identity could be decoded within each session a within condition decoding analysis was performed for each session with an SVM (see above), using the MATLAB function *fitcecoc* and 10-fold Cross-validation.

##### Decoding task condition

We performed binary decoding with a SVM (imagery trial = 0, viewing trial = 1) as a function of number of neurons. For each population size *k* = [1: *N*_*neurons*_] we randomly selected *k* neurons without replacement and trained a SVM with 10-fold cross validation. To ensure reliability the entire sampling and decoding procedure was repeated 100 times for each population size leading to a distribution of accuracy values. Even with just one neuron, it was possible to distinguish between task states (Figure 5F).

#### Object ‘reconstruction’ during imagery

Reconstructions of objects during the imagery period were carried out in a manner similar to object reconstruction during viewing (Figure 3D). For all sets of simultaneously recorded, axis-tuned, and reactivated neurons (neurons in all sessions that satisfied all three criteria), we learnt a linear mapping between the stimulus features and the firing rates during imagery and decoded the feature vectors for all images out of sample using leave-one-out cross validation. Given the relatively low number of neurons in each session that satisfied all three criteria only the first five feature dimensions were used instead of the full fifty, so that the number of neurons exceeded the number of dimensions needing decoding. The reconstructed images have high similarity upon visual inspection (Figure 5E), and the normalized distances in visual feature space are closer than chance (Figure 5D, median = 0.89). Note that for 64/94 stimuli that were reconstructed, the best possible reconstruction was of a different image type (i.e. different semantic identity).

#### Imagery Selectivity Index

To quantify the relative preference of each of the 231 neurons for imagery vs. viewing (encoding) we computed an imagery selectivity index. Mean firing rates across all trials (regardless of image identity) were calculated over the full post-stimulus periods (5 s for imagery, 2 s for encoding). The imagery selectivity index was defined as:

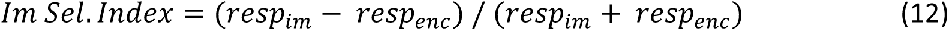

Where *resp*_*im*_and *resp*_*enc*_represent the mean firing rates during imagery and encoding periods, respectively. This index ranges from -1 (encoding selective) to +1 (imagery selective), with values near 0 indicating similar mean responses to both conditions.

#### Imagery Heatmap Construction

Neurons were rank-ordered by their imagery selectivity index in descending order (most imagery-selective to most viewing-selective). Binned mean responses were computed in a sliding window. For imagery, the bin size was 1000 ms with 100 ms step size starting at 500 ms before the onset of imagery. For encoding, the bin size was 400 ms bin with 40 ms step size starting at 200 ms before image onset. The two (mean imagery response and mean encoding response) were concatenated so that each row represented a single neuron and columns represented sequential time bins. The responses were then z-scored across rows and plotted as a heatmap (Figure 5F).

## Acknowledgements

We thank the staff of the epilepsy monitoring unit (neuromonitoring staff, nursing staff, and physicians) and of the Biomedical Imaging Research Institute at Cedars-Sinai Medical Center for patient care and support. We thank Emily Choe for help implementing the GAN used in Figure 3. We thank members of the Tsao and Rutishauser labs namely Janis Hesse, Liang She, Le Chang, Pinglei Bao, Yuelin Shi, Hristos Courellis, and Austin Brotman for helpful comments throughout all stages of this project. We thank the patients for all their patience and perseverance. We thank the three anonymous reviewers for helping us to greatly improve the content and clarity of our manuscript.

## Author contributions

V.W., U.R., and D.Y.T designed the study. V.W. collected and analyzed the data. V.W., U.R., and D.Y.T. wrote the paper with input from C.M.R, J.M.C., L.M.B., and A. N. M. C.M.R, J.M.C., and L.M.B. provided patient care and facilitated experiments. A. N. M. performed surgery.

## Data and materials availability

Data and code will be made publicly available upon acceptance.

## Supplementary materials

## Supplementary results

### Axis code is independent of the specific deep network used to parametrize stimuli (relevant for Figures S3 & S4)

All analyses reported in the main text use a feature space built from layer fc6 of AlexNet. However, there are many other deep convolutional neural network models that achieve high performance on object recognition (*108*). We therefore set out to compare the ability of several such models to explain the responses of VTC neurons.

The models tested include AlexNet (*109*), both VGG-16 and 19 (*109*), VGG-Face (*110*), the eigen object model wherein the space is built by performing PCA on the pixel level representation directly (*111*, *112*), four CORNet models (*113*, *114*) and an unsupervised Vision Transformer model (DINO-ViT) (*72*). AlexNet, VGG-16/19, and the CORNet family of networks are trained to classify images into 1000 object categories with varying architectures. AlexNet has 8 layers, with 5 convolutional and 3 fully connected layers. VGG-16 and 19 have 16 and 19 layers respectively, with 3 fully connected layers and the rest being convolutional layers. The CORNet family of networks consists of 4 networks: CORNet-Z is purely feedforward; CORNet-R/RT includes some recurrence which has been shown to be essential for object recognition in the primate visual system (*115*); and finally CORNet-S has the most complicated of architecture, including recurrent and skip connections between the layers (*114*). Despite the individual differences all four networks have architectures inspired by the primate visual system with layers corresponding to V1, V2, V4, and IT. VGG-Face has the same architecture as VGG-16 but is trained to identify 2622 celebrities (*110*). DINO-ViT is a Vision Transformer model trained using self-supervised learning, without using labels. ViT was trained on a curated dataset (LVD-142M) of 142 million images (*72*).

To quantify the ability of each network to explain human VTC neural responses we learned a linear mapping between the features of each model and the neural responses (*116*). We reduced dimensionality of the feature representations via principal-component analysis (PCA) yielding 50 features for each object and model. In our main analyses, we used leave-one-out cross validation and for each neuron fit the responses of 499 images to the 50 features via linear regression before predicting the response of the neuron to the one left-out image using the same linear transform. The amount of variance explained of the label of the left-out image by the predicted response was used as the measure of goodness-of-fit (Figure S3C). We also computed the encoding and decoding error for each neuron for every model. Encoding error (Figure S3E) was computed as follows: the observed population vector to each object was compared to the predicted population response vector and the observed population response vector to a random other object in the set. If the angle between the observed and predicted response to the chosen object was smaller than the angle between the predicted response and the distractor, the prediction was considered correct (Figure S3D). Decoding error (Figure S3F) was computed via the same method except the feature vector of the object was predicted from the neural responses. In other words, the roles of the neural responses and the model features were reversed (see supplementary methods). A model was considered to explain neural responses well if it has high explained variance and low encoding/decoding error. The worst performing models were the eigen object model and VGG-Face (Figure S3C, E, F), while the best performers were DINO-ViT and the most complicated CORNet (CORNet-S). DINO-ViT explained significantly more variance than all other models except CORNet-S (p = 0.095, Friedman test with Tukey-Kramer post-hoc correction). All remaining models, including AlexNet, performed similarly well. (Figure S3A; p < 0.001, AlexNet vs VGG-Face; p < 0.001, AlexNet vs eigen model, Friedman test with Tukey-Kramer post-hoc correction).

### A note on the heterogeneity of mental representations across people (relevant for Figure S7D)

There is a large body of evidence to support the notion that the subjective vividness of visual imagery varies greatly between individuals (*37*, *117*, *118*), with some individuals demonstrating a complete inability to generate a mental image (aphantasia) (*119*) while others have near-photorealistic mental images (hyperphantasia). Moreover, neuroimaging studies have shown differences in fMRI BOLD signals — both intensity in early visual areas (*120*) and functional connectivity between areas (*117*, *121*) — between subjects that reported different amounts of vividness in imagery. Growing evidence of these differences has led to the conclusion that examining mental imagery at the group level with current tools (fMRI and psychophysics) is not appropriate — leading to the end of the “imagery debate” (*122*, *123*).

In order to understand whether there is a correlation between the data discussed here and subjective vividness, patients completed the Vividness of Visual Imagery Questionnaire (VVIQ) (*124*). All patients we recorded from had relatively high scores in the vividness scale. For comparison, the median VVIQ value in a recent international survey of 9063 participants was 58 (*125*), with 89.7% (8129/9063) falling within the category of ‘typical visual imagery’. In our dataset, 5/6 patients fell within the ‘typical visual imagery’ category, and one into the ‘hyperphantasic’ category (Figure S7D). An important open question that remains is whether the recapitulation of sensory context we found is weak or absent in aphantasic individuals.

## Supplementary Methods

### Model comparisons

#### Extraction of visual features from stimulus images

Each stimulus image was fed into one of the following models to extract the corresponding features:

#### Eigen object model

PCA was performed on the original images of the 500 stimulus objects and the top 50 PCs were extracted to compare with other models.

#### AlexNet

We used the pre-trained MATLAB implementation of AlexNet. This is an 8 layer deep convolutional neural network with 5 convolutional layers and 3 fully connected layers, trained to classify images into 1000 object categories.

#### VGG Family

We used the pre-trained MATLAB implementations of VGG-16, a 16 layer deep convolutional neural network that contains 16 layers with 13 convolutional layers and 3 fully connected layers trained to classify images into 1000 object categories (*109*), VGG-Face which has the same structure as VGG-16 but is trained to recognize the faces of 2622 celebrities (*110*), and VGG-19 which has 19 layers (16 convolutional and 3 fully connected) trained on the same task as VGG-16 (*109*).

#### CORNet Family

We used a pre-trained PyTorch implementation of CORNet. The CORNet family contains 3 architectures: CORNet-Z, CORNet-R/RT, and CORNet-S. Each architecture includes 4 main layers that correspond to V1, V2, V4, and VTC. CORNet-Z is the simplest model and is purely feedforward. CORNet-R/RT takes the otherwise feedforward network and introduces recurrent dynamics within each area. CORNet-S is the most complex containing within area recurrent connections, skip connections and the most convolutional layers.

The parameters of AlexNet and VGG were obtained from MATLAB’s Deep Learning Toolbox. The CORNets were downloaded from (https://github.com/dicarlolab/CORnet).

#### DINO-ViT

We used a pre-trained Vision Transformer (ViT) model trained using self-supervised learning without using any labeled data. The model employs a student-teacher distillation framework wherein a large (ViT) with 1 billion parameters serves as the teacher, which is then distilled into smaller models (ViT-Small, ViT-Base, and ViT-Large; ViT-Base was used for our analysis). The model was trained on 142 million curated images sampled from other known image sets such as ImageNet-22k, the training set of ImageNet-1k, and Google Landmarks amongst others (see Table 15 in (*72*)) rather than a single labeled dataset. Through its self-supervised training approach, the model learns visual representations that capture fundamental image features— such as object boundaries, textures, spatial relationships, and semantic content—that generalize across multiple vision tasks including classification, segmentation, and depth estimation, without requiring task-specific fine-tuning. The model was downloaded from (https://github.com/facebookresearch/dinov2).

#### Extraction of semantic features

Semantic features for all the images shown to patients were obtained in a multistage process. First, ImageNet labels for all images were automatically obtained and manually reviewed and replaced with the correct labels from the ImageNet taxonomy if needed. Second, vector embeddings for the resulting ImageNet labels were obtained for eight distinct word embedding spaces (see below) that differed in training methodology, corpus size, and dimensionality. For single word labels, vectors were directly retrieved from the pre-trained embedding space using the model’s native lookup functionality. For two-word labels (for example, ‘coffee machine’) the embedding vector was obtained by averaging the individual word vectors.

#### GloVe (Global vectors for word representation) family

We used four pre-trained variants of GloVe (*126*), all trained using global matrix factorization. GloVe 6B.300d is trained on Wikipedia 2014 and Gigaword 5 corpora with 6 billion tokens and has a 400k word vocabulary. GloVe 42B.300d and GloVe 840B.300d are trained on Common Crawl web corpus with 42 billion tokens and 840 billion tokens respectively. The former has a 1.9M word vocabulary, the latter has a 2.2M word vocabulary. GloVe Twitter is trained on the Twitter corpus (2 billion tweets, 27 billion tokens), it has a 1.2M word vocabulary that is more colloquial than the others. The embedding spaces of the first three models have 300 dimensions, and the twitter space has 200 dimensions. All variants were downloaded from (https://github.com/stanfordnlp/GloVe).

#### Google News Word2Vec

We used a pre-trained word2vec (*126*) that produced 300D vectors, and was trained on the Google News corpus using a skip-gram model with negative sampling. It was trained with 100 billion tokens and has a 3M word vocabulary. The model was downloaded from (https://github.com/mmihaltz/word2vec-GoogleNews-vectors).

#### FastText

We used three FastText word2vec variants (*127*) that used continuous-bag-of-words (CBOW) with position-weights, character n-grams of length 5, window size 5, and 10 negatives. We used MATLAB’s built in pre-trained embeddings (accessed via the *fastTextWordEmbedding* function), which was trained on the Wikipedia, UMBC web base, and statmt.org news corpora with 16 billion tokens and a 1M word vocabulary. CC.en.300.vec which is part of the multilingual FastText collection (*128*) and trained on the Wikipedia and Common Crawl corpora. Crawl-300d-2M.vec which is trained on the Common Crawl corpus with 600 billion tokens and has a 2M word vocabulary (*129*). All FastText embeddings were 300D.

All embeddings were reduced to 50 dimensions via PCA to be consistent with the axis model. 50 dimensions explained ∼70% of the feature variance in 300D models and ∼80% of the feature variance in the 200D model.

#### Category models with auto-clustering

The category model used in the main text is a generalized linear model (GLM) with a Poisson response variable and five binary pre-defined categorical indicators. We also carried out a control analysis, as part of which we assigned category labels automatically using clustering. This way, we could systematically compare different category models with different numbers of categories (clusters) to assess which explains best VTC responses. We used a range of *N*_*cats*_ = [1: 25] number of categories, and compared the neural variance explained of each to the axis model. We compared the following different ways of assigning images to categories:

##### Hierarchical clustering based on the neural response

An initial similarity structure was established by computing the Pearson correlation coefficients between all stimulus pairs based on their neural response across the population. This correlation matrix was converted to a dissimilarity matrix (D = 1 – correlation) for hierarchical clustering using average linkage. Cluster assignments for a chosen *N*_*cats*_were determined by cutting the dendrogram at a varying number of levels.

##### Visual feature clustering

Visual features of the stimulus images were extracted from all models described above (AlexNet, VGG family, CORNet family, and eigen model) and clustered in 50D object space using k-means clustering.

##### Semantic feature clustering

Semantic features for all stimulus images were obtained from all models as described above (GloVe family, Google News Word2Vec, and the FastText family) and clustered in 50D semantic space using k-means clustering.

Explained variance was computed for each categorical representation (*N*_*cats*_ = [1: 25], semantic, visual) and compared to the explained variance of the axis model with 25 dimensions. The axis model with 25 dimensions gave the best performance for fc6 (Figure S2G)

### Explained variance computation

See main methods (‘Axis model’ subheading). The reported results are the ratio of explained to explainable variance in the 318/367 axis-tuned neurons that had a > 10% explainable variance.

### Encoding/Decoding error computation

For the encoding analysis, the response of each neuron was first z-scored (with the mean and standard deviation computed based on the response), then the same procedure as the explained variance computation was followed to obtain a predicted response to every single object. To quantify prediction accuracy, we examined the angle between the predicted population response to each object and its actual population response (target) or the population response to a different object (distractor). If the angle between the predicted response vector and one of the distractors was smaller than the angle between the predicted response vector and the target this was counted as an error. Overall encoding error was quantified as the average errors across 1000 pairs of target and distractor objects.

For the decoding analysis we used exactly the same procedure, however the roles of the neural responses and the object features were reversed. We first normalized each dimension of object features to have zero mean and unit variance, then for each image we used a leave-one-out procedure to fit a linear transform using the responses to 499 images and then predict the features of the left-out image. Decoding error was computed as the average decoding error across all target and distractor pairs in feature space.

### Quantification of axis consistency

The consistency of the preferred axis of a neuron (Figure S2E) was determined as follows: The image set was randomly split into two subsets of 796 and 797 images, a preferred axis was computed for each set and the Pearson correlation was computed between the two. This procedure was repeated 100 times and the axis consistency was defined as the to get an aggregate correlation value.

## Supplementary figures

**Figure S1.**
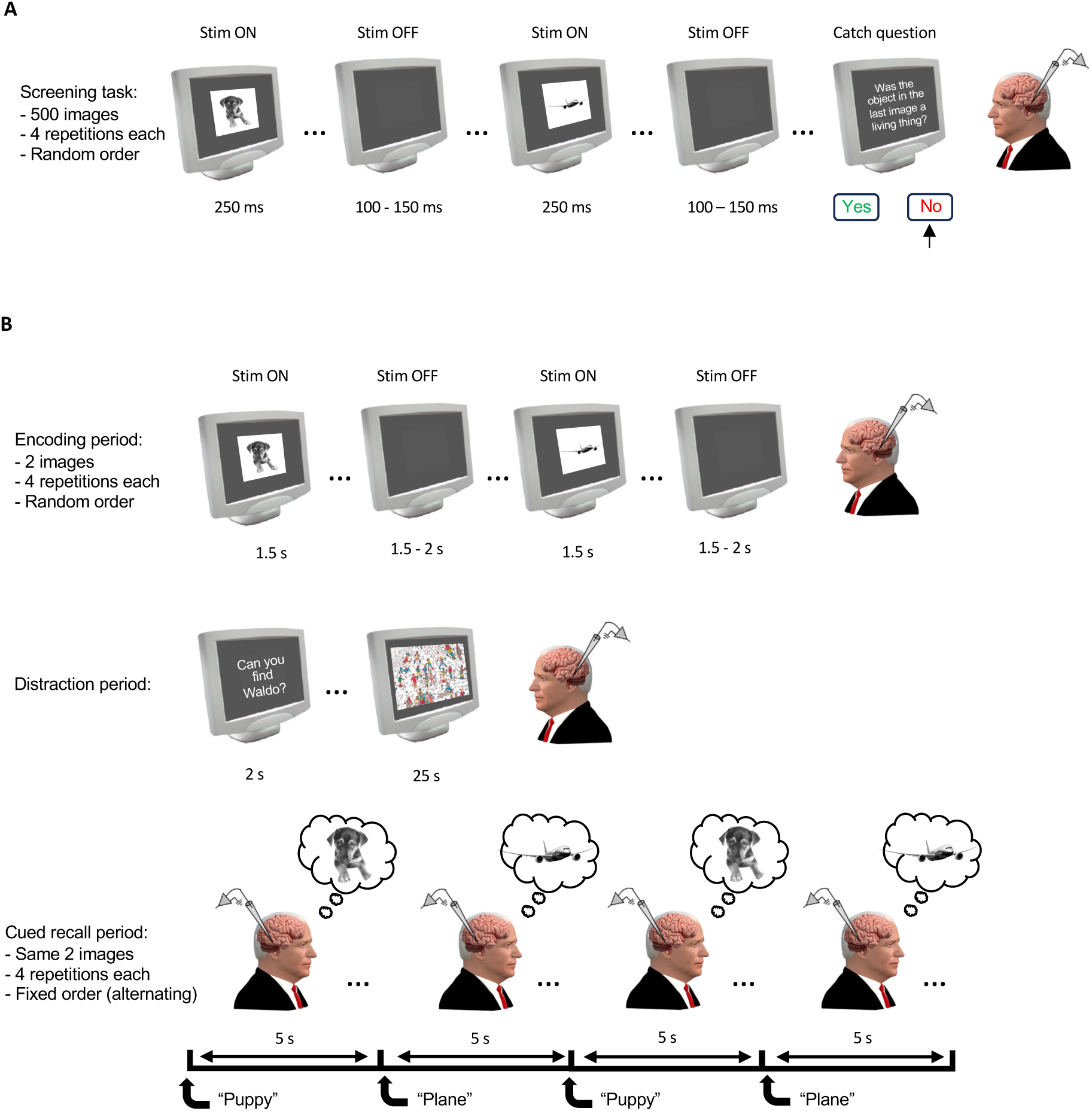
Detailed schematics of tasks. **(A)** Screening task. Grayscale images with white backgrounds were displayed on a gray screen for 250 ms with the inter-trial interval jittered between 100-150 ms. Images subtended 6-7 visual degrees. At random intervals (min interval: 1 trial, max interval: 80 trials) a yes-no catch question appeared pertaining to the image that came just before it. Each image was repeated 4 times for a total of 2000 trials **(B)** Imagery task. A subset (6–8) of the 500 images used for screening were used, chosen to have spread across both the preferred and principal orthogonal axes (see Figure 4A, B for examples of images chosen). Two images were used per trial. Each trial consisted of an initial encoding period wherein the two images were displayed one at a time on a gray screen for 1.5 s with the inter trial interval jittered between 1.5 – 2 s. After each image was viewed 4 times, a distraction period occurred wherein patients were required to spend 30 s on a visual search puzzle. Finally, after the distraction period they were cued verbally by the experimenter to imagine both images in the trial one by one in alternating order until both had been visualized 4 times. Each image appeared in 2 trials for a total of 8 repetitions of viewing and imagery per image.

**Figure S2.**
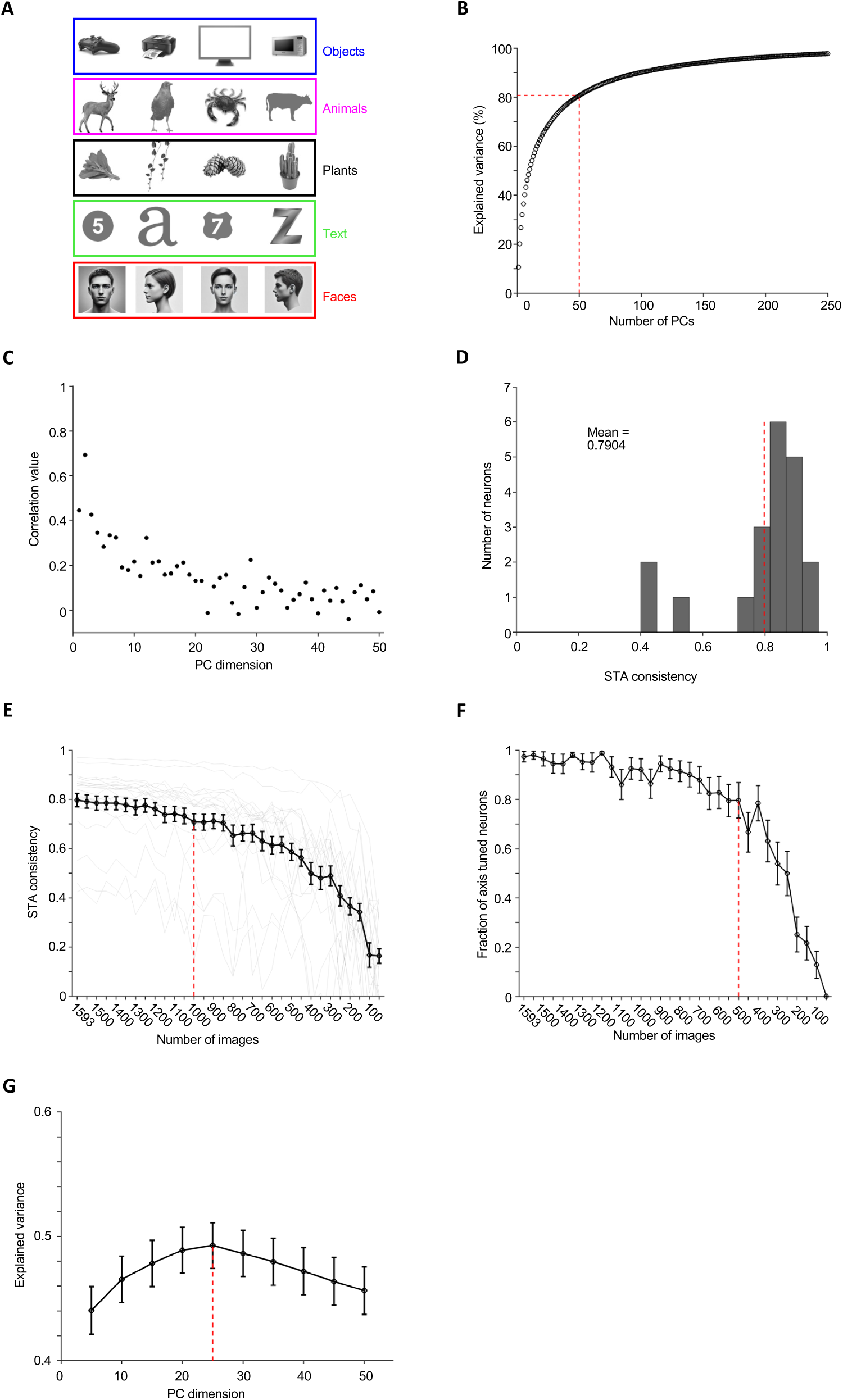
Stimuli and parameters chosen for screening task. [Note: The human faces in panel A of this figure have been replaced with synthetic faces generated by a diffusion model (*130*), in accordance with the bioarxiv policy on displaying human faces.] **(A)** 500 stimuli from the categories face, text, plant, animal, and object were shown to the patients. Panel shows example images from each of the 5 categories (taken from www.freepngs.com). **(B)** The cumulative explained variance of fc6 unit as a function of the number of principal components used. 50 dimensions explained 80.78% of the variance. **(C)** Correlation between predicted feature values using the axes computed via the 500 stimulus screening and actual feature values for all 50 dimensions. In Figure 3B we plot the predicted and actual feature (fc6) values for all images and compute the correlation for PC1. This plot extends that analysis to all 50 PCs. The fc6 features are predicted using a leave-one-out paradigm wherein we use the responses to all but one object (500-1 = 499) to determine the transformation between responses and feature values by linear regression, and used that transformation to predict the feature values of the held-out object. **(D-F)** In one early session a patient performed a more comprehensive version of the screening task, with 1593 stimuli (*56*). The 500 used in this project were subsampled from this larger set. The responses of the 22 axis-tuned neurons in this session were used to determine appropriate parameters going forward. **(D)** Distribution of axis consistency (computed using half-splits, see supplementary methods) for axis-tuned neurons in this session. Red line indicates the mean value. **(E)** Axis consistency as a function of the number of stimuli used to compute axes. The red line at 1000 indicates how consistent axes would be if computed using 500 stimuli. Axis consistency remains stable until the stimulus number drops below 400 in each half split. This informed the choice of 500 stimuli. **(F)** Proportion of axis-tuned neurons detected as a function of the number of stimuli used. 78% of the axis-tuned neurons detected using 1593 stimuli were still detected using 500 stimuli. At lower stimulus numbers the neuron count drops precipitously. **(G)** Explained variance as a function of the dimensionality (number of PCs) of object space. We fit axes to neural responses using a varying number of dimensions and tested for their predictive accuracy on responses to held out images (see methods). The best performing axis model uses 25 dimensions rather than all 50.

**Figure S3.**
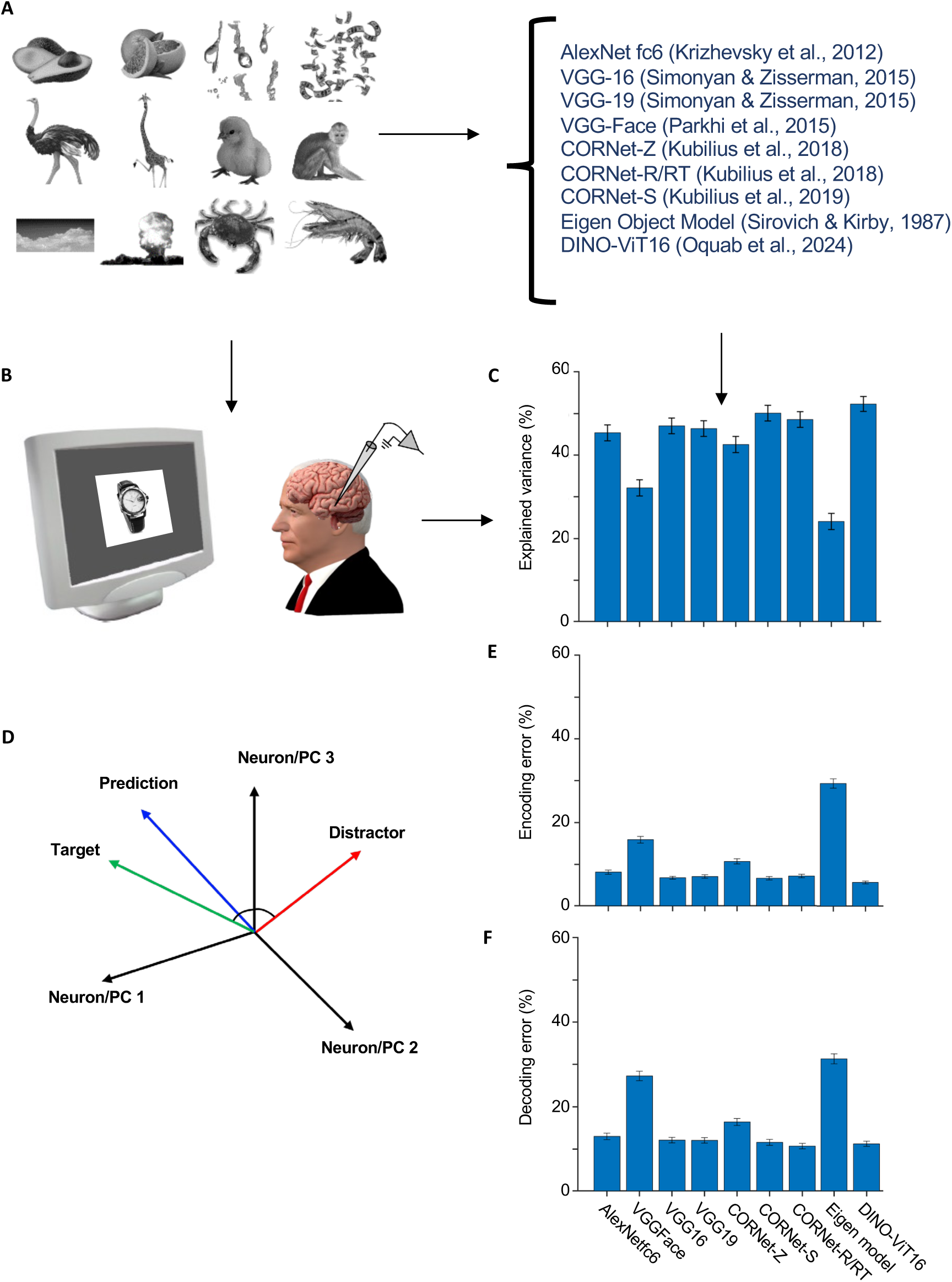
Comparison of different models. **(A)** The 500 grayscale stimulus images used in the screening task were parametrized using 9 different models from 5 different model families (AlexNet, the eigen object model, VGG, CORNet, and a vision transformer). The same number of features were extracted from units of the different models using principal components analysis (PCA) for comparison. **(B)** Responses from VTC neurons were recorded as patients viewed these objects 4 times each. For recording locations and task schematic see Figure 1D and Figure S1A respectively. **(C)** The explained variances for each model after 50 features were extracted using PCA. For each neuron, explained variance was normalized by the explainable variance (see methods). Error bars represent SEM for the recorded neurons. **(D-F)** The different models were compared with respect to how well they could predict the neuronal responses or the object features. In both cases a leave-one-out procedure was used to learn and test the transformation between responses and features. To quantify encoding error for example, for each object we compared predicted responses to individual objects in the neural state space to the actual responses to that object and a distractor object. **(D)** If the angle between the predicted response and the actual response was smaller than the angle between the predicted response and the distractor the encoding was considered correct. To quantify decoding error, we reversed the roles of the neural responses and the object features and decoded object features before comparing the decoded features to the actual features for a given object and a distractor object. **(E)** Encoding error across all models. **(F)** Decoding error across all models. The eigen model, VGG-Face, and the purely feedforward CORNet-Z had larger encoding/decoding errors than the other models, consistent with them explaining less variance as well.

**Figure S4.**
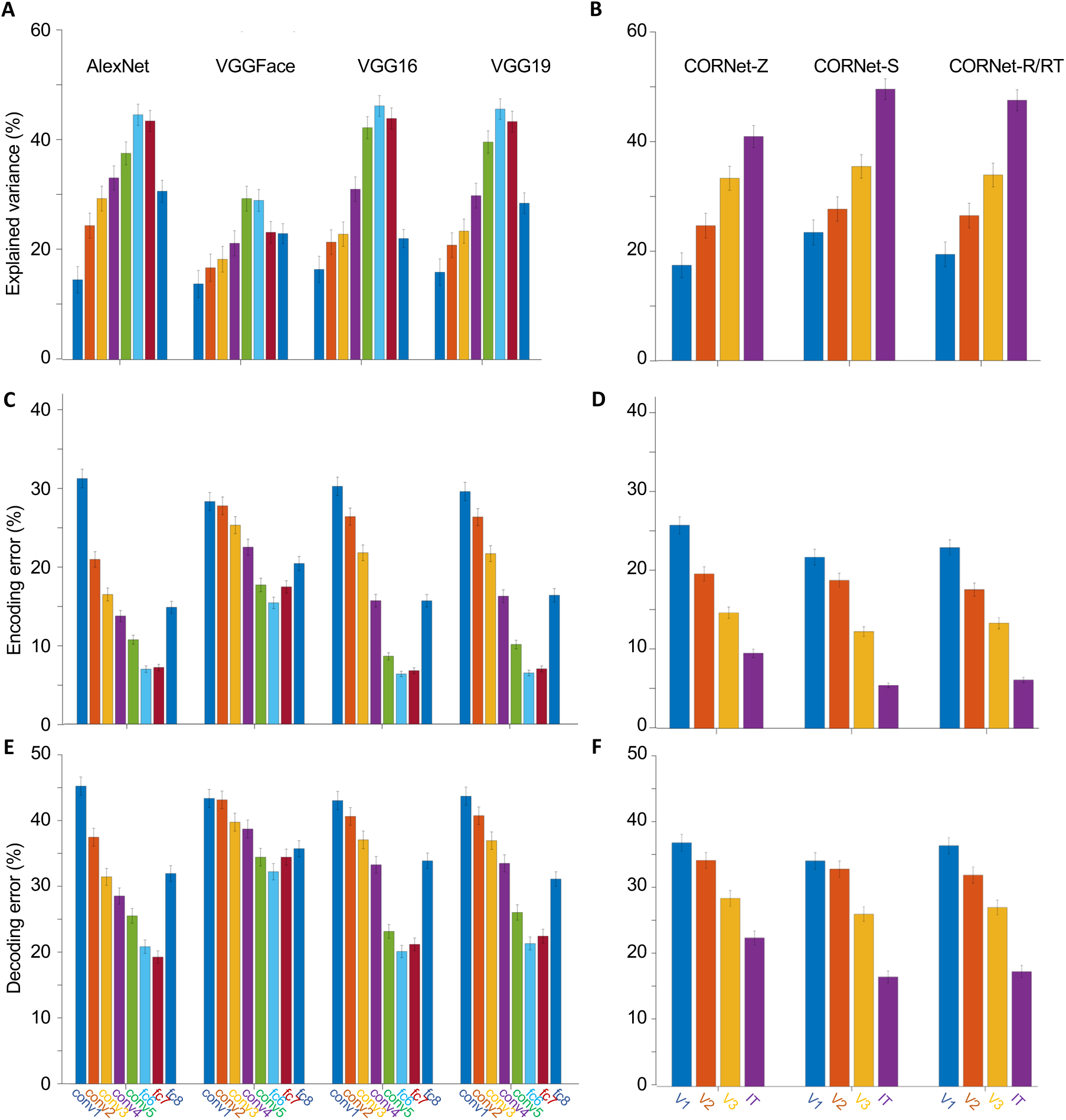
Comparing explained variance, encoding, and decoding error across layers of each model. Inspired by previous work (*56*, *68*, *131*) units in the penultimate layer were used to build the object space used in all analyses. Here we compare performance across all fully-connected layers of AlexNet and the VGG networks, and all output layers of the CORNet networks used in Figure S3. **(A)** Explained variance for the main layers of AlexNet and VGG models. **(B)** Explained variance across the layers of the various CORNet models (V1, V2, V4, IT). The ‘IT’ or VTC layer performs the best in all CORNet versions, with little difference across the various forms with the exception of the purely feedforward version (CORNet-Z) performing worse than its recurrent counterparts. **(C)** Encoding error across layers of AlexNet and VGG models. As expected, the encoding error is lowest for the fully connected layers with the highest explained variance. **(D)** Encoding error across layers of the CORNets. **(E)** Decoding error across layers of AlexNet and VGG networks. **(F)** Decoding error across layers of CORNets.

**Figure S5.**
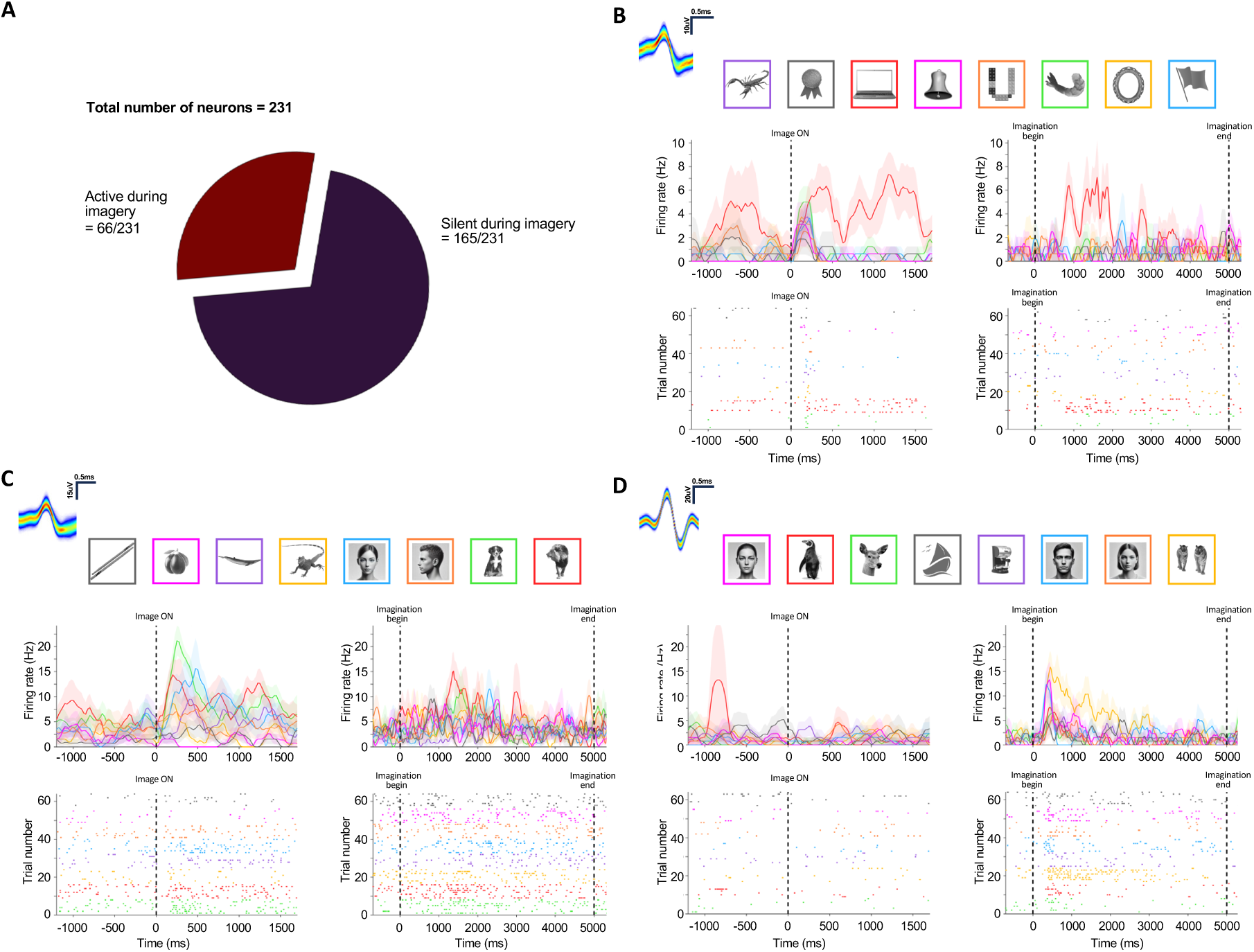
Reactivation of VTC neurons during imagery. [Note: The human faces in panel B-D of this figure have been replaced with synthetic faces generated by a diffusion model (*130*), in accordance with the bioarxiv policy on displaying human faces.] **(A)** The proportion of total recorded VTC neurons that reactivated during imagery. **(B-D)** Example neurons. Left PSTH and raster shows the response of the neuron during the encoding while right PSTH and raster shows the response during imagery. The stimulus images used in the task are shown above arranged in ascending order of projection onto the neurons preferred axis (computed during screening). **(B)** This neuron’s preferred stimulus was the laptop during encoding and it reactivated robustly during imagery of the laptop. Note that this was not an axis-tuned neuron and as such the laptop is not the right most image. **(C)** Example of an axis-tuned unit that reactivated to multiple stimuli (red & green) in a graded manner during imagery. **(D)** A small number of neurons recorded (15/231) were quiet during encoding but strongly active during imagery. Once such example is shown.

**Figure S6.**
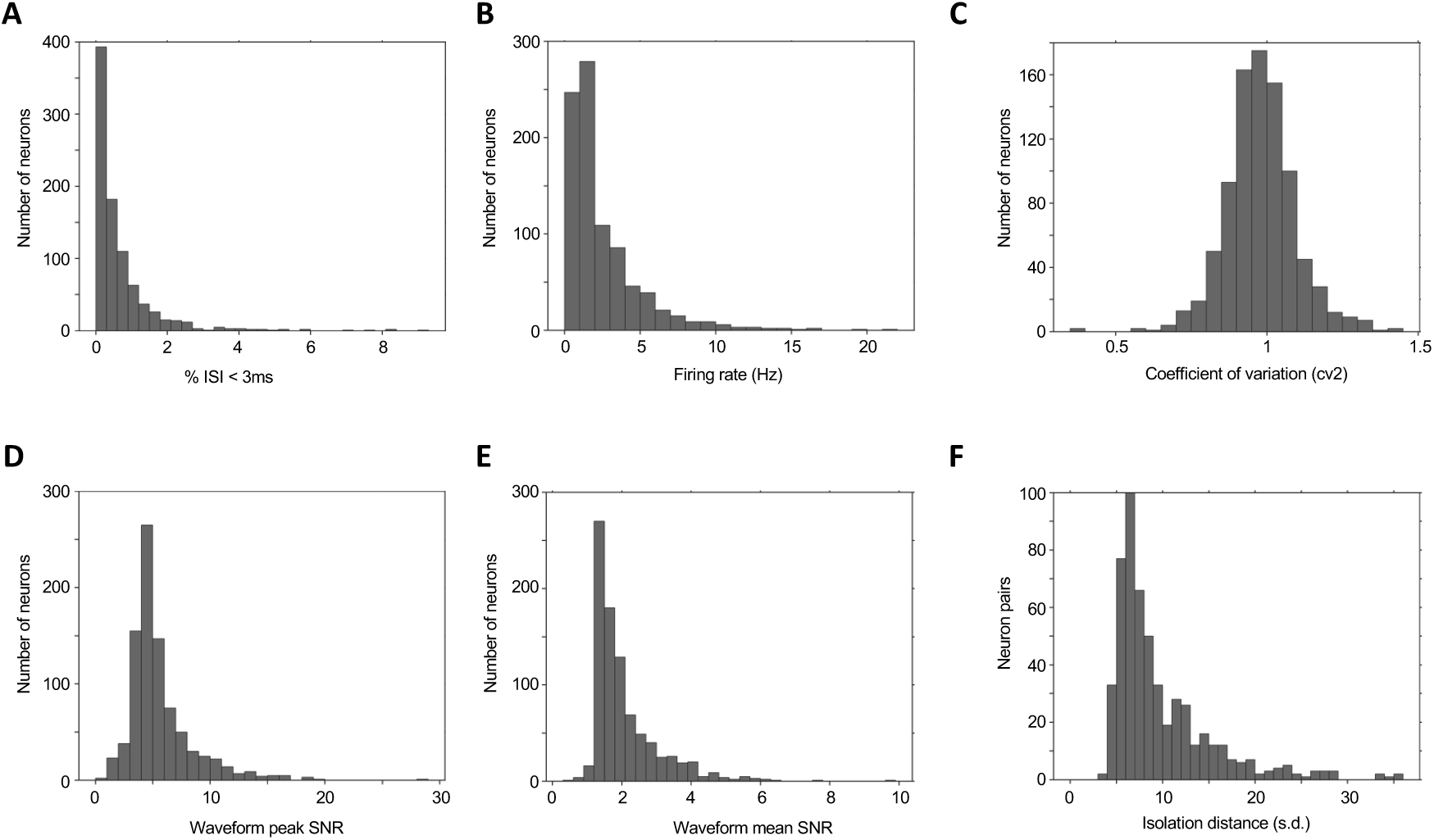
Spike quality metrics for all identified putative single units. **(A)** Proportion of inter-spike intervals (ISI) below 3 ms. **(B)** Average firing rate. **(C)** Coefficient-of-variation. **(D)** Signal-to-noise ratio (SNR) for the peak of the mean waveform across all spikes as compared to the standard deviation of the background noise. **(E)** Mean SNR of the waveform. **(F)** Pairwise distance between all pairs of neurons on channels where more than one neuron was isolated.

**Figure S7.**
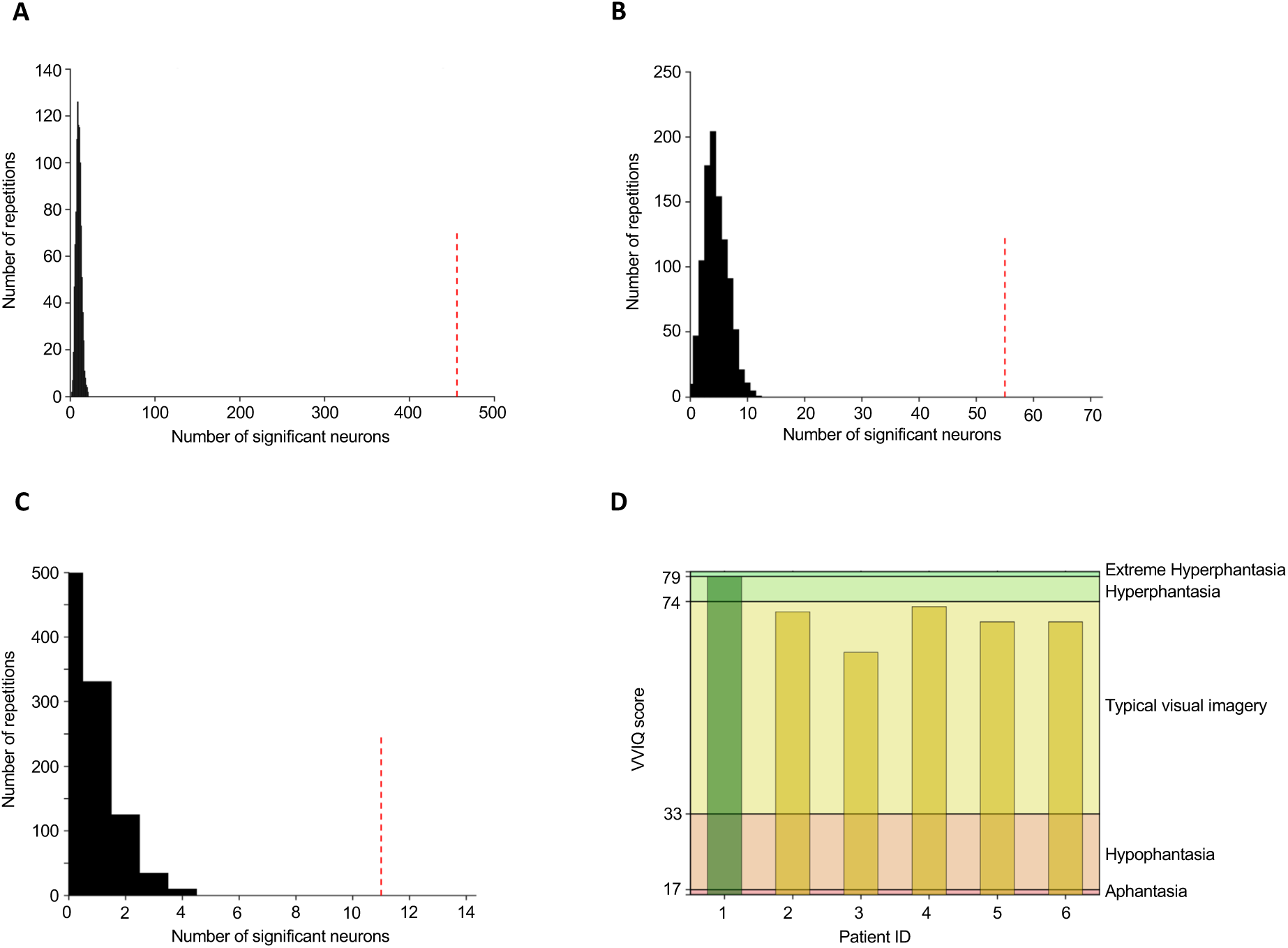
Permutation statistics for number of neurons selected as visually responsive and active during imagery. **(A)** Significance of the number of visually responsive VTC neurons. The dashed red line indicates the number of neurons selected as visually responsive (456/714). The null distribution (black) was estimated by re-running the identical selection procedure (sliding window ANOVA, see methods) after randomly shufling the trial labels. Shufling the trial labels destroys the association between the spiking response and trial identity, but keeps everything else (trial number, stim ON time etc.) intact. The shufling procedure was carried out for 1000 repetitions. The mean of the null distribution was 16, implying that the chance level for selecting a neuron as visually responsive is 16/714 or ∼2%. The p-value reported is the percentage of null distribution values that are greater than the chosen number of neurons. In this case, p = 0 and is reported as 1/number of repetitions p = 0.001. **(B, C)** Significance of the number of neurons in VTC active during imagery. Given that the selection criteria for activation during imagery is either a number of consecutive significant bins of a sliding window ANOVA or sliding window t-test we computed a null distribution for each. The null distribution for each is computed by re-running the identical selection procedure for 1000 repetitions after randomly shufling the spike times for the t-test and the trial labels for the ANOVA in the visual responsivity test (see A). The mean of the null distribution for the t-test (B) was ∼5, implying that the chance level for labeling a neuron as active during imagery via t-test is 5/231 or ∼2% and p = 0.001, while for the ANOVA the mean is ∼1 so the chance level is 1/231 or ∼0.5% and p = 0.001. **(D)** ‘Vividness of visual imagery’ questionnaire scores of patients that performed the cued imagery task. All patients recorded from in this study scored highly in the VVIQ or having good visualization capabilities.

**Figure S8:**
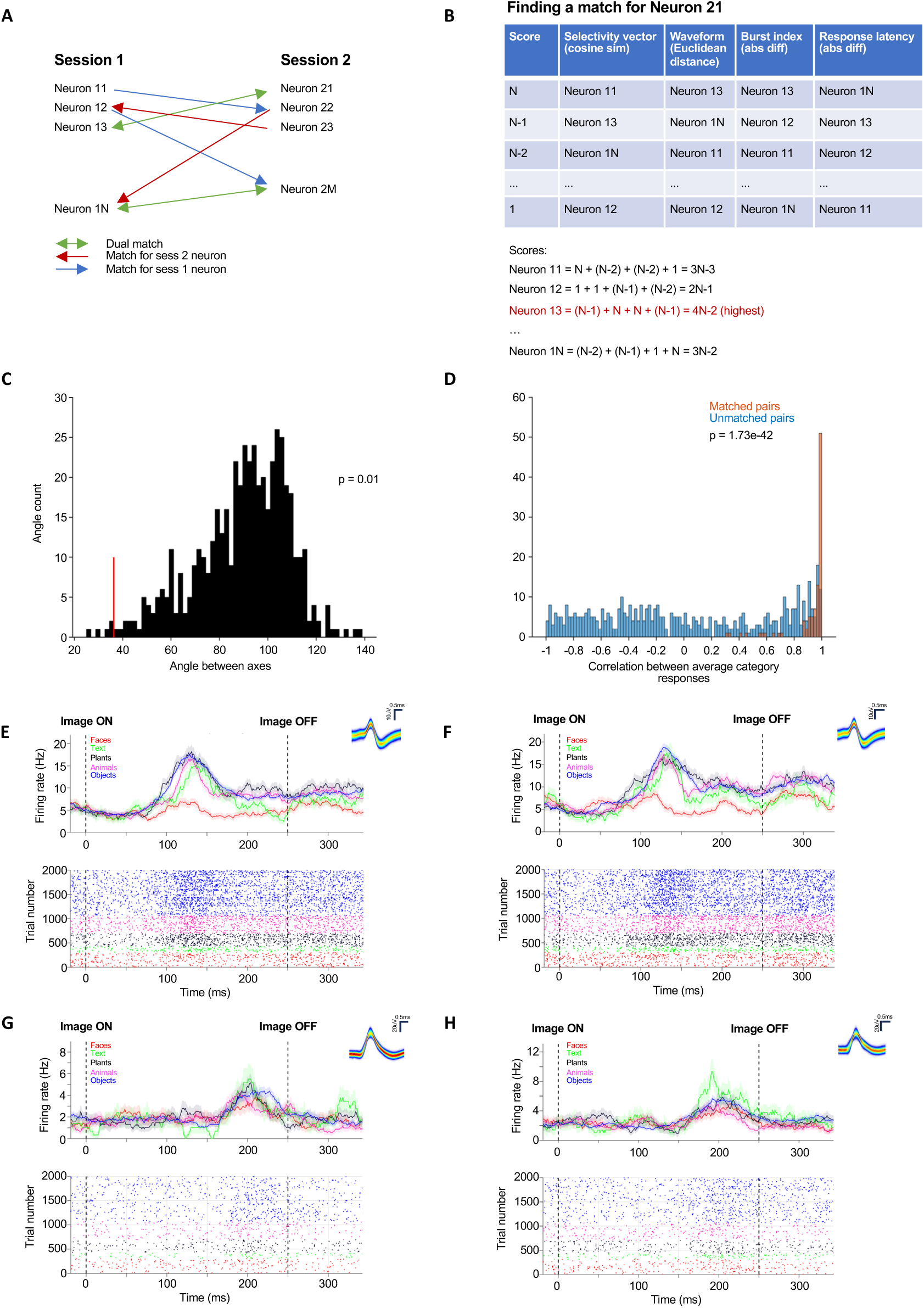
Matching neurons across sessions. **(A)** Schematic of matching procedure. Matching of neurons was done in a bi-directional manner. Each neuron in the first session was evaluated for similarity against all neurons in the second session using the waveform, selectivity, angle of axes (if axis-tuned), response latency, channel number, and burst index (see methods) while keeping track of its closest counterpart. Then each neuron in the second session was compared against all neurons in the first session. If a pair of neurons were both chosen as each other’s closest counterpart, then the match was automatic. If not, the match had to be made manually. All matches were manually verified by looking at the shape of the peri-stimulus time histogram (PSTH) at the end. **(B)** Example: Computing matching score for example neuron N21 (session 2, neuron 1). The rank orders were assigned scores (rank 1 = n points and rank n = 1 point, where n = number of neurons in session one being compared) for each metric and then totaled. The session two neuron with the highest match score (neuron N13) is marked as a match. In this example, as N21 was initially chosen as the closest match for N13 amongst session 2 neurons, this match is automatic. **(C-D)** Metrics to evaluate matching quality. **(C)** Angles between axis of pairs of neurons. Well matched pairs are expected to have small angels. Red line indicates the average angle between matched neurons (37 degrees), black distribution shows angles between pairs of non-matched axis tuned neurons (mean = 86 degrees). The average angle between matched neurons is significantly smaller than the angles between non-matched neurons (p = 0.01, comparison to null distribution). **(D)** Correlation of mean category response between matched (orange, mean value = 0.90) and non-matched (blue, mean value = 0.02) pairs of neurons. The distribution of matched neuron values is significantly larger (p = 1.73e-42, Kolmogorov-Smirnoff test). **(E-F)** Examples of automatically matched neurons. **(E)** Neuron from session one that was most closely matched to the neuron in (F). **(F)** Neuron from session two most closely matched to the neuron in (E). **(G-H)** Example pair of neurons that were matched manually, as the automatic matching was not bijective.

**Figure S9:**
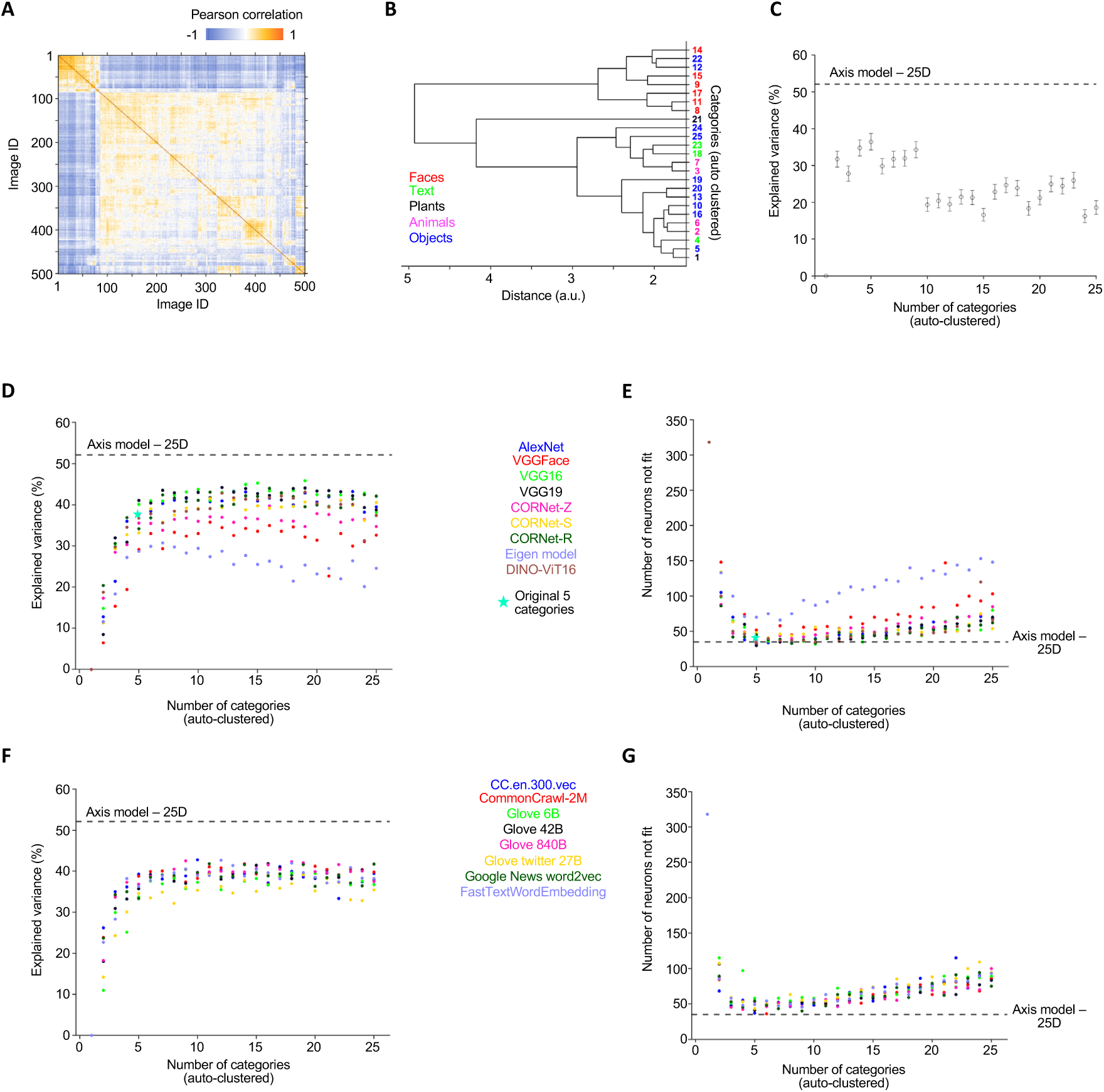
Comparison of neural variance explained between the axis model and different category models. We examined (A-C) Hierarchical clustering based on the neural response, (D-E) Clustering of visual features, and (F-G) clustering of semantic features. **(A-C)** Hierarchical clustering of categories based on neural response. **(A)** Pairwise similarity matrix of the neural responses of all 500 images (Pearson correlation coefficients). The images are sorted as shown in (B). **(B)** Dendrogram built from the similarity matrix in (A) with average linkage and up to 25 clusters (categories). The cluster numbers are listed on the y axis and colored by the originally used 5 categories. **(C)** Explained variance as a function of number of clusters. The clusters are chosen by cutting the dendrogram in (B) off at different points. The dashed line indicates the explained variance of the axis model in fc6 with 25 dimensions (best performing axis model, see Figure S2G), which was significantly higher than clustering with any of the tested number of categories (p = 3.60e-20, paired t-test between axis model and best performing number of clusters *N*_*cats*_= 5). **(D-E)** Creating and evaluating visual categories by feature clustering. Visual features of the stimulus images were extracted from the models indicated, followed by clustering in 50D object space using k-means clustering. For each neuron, a category model with the images assigned to each of the identified cluster was then fit to evaluate the amount of explained variance. **(D)** Neural explained variance as a function of number of categories and model type. The dashed line indicates the explained variance of the axis model (fc6 with 25 dimensions) which was significantly higher (p = 1.64e-10, paired t-test between axis model and highest performing clustering; VGG16 with 19 clusters). The performance of the category model with the pre-defined 5 categories is indicated with a star. **(E)** Number of neurons for which a given model did not explain a significant amount of neural variance (explained variance < 0). The lower the number of neurons not fit, the better the model. The 5-category model with the pre-defined 5 categories is indicated with a star. **(F-G)** Same as (D-E), but for semantic models. Semantic features for all images were extracted by first obtaining ImageNet labels for all images. **(F)** Neural explained variance as a function of number of categories and model type. The dashed line indicates the explained variance of the axis model in fc6 with 25 dimensions which was significantly higher (p = 1.41e-12, paired t-test between axis model and highest performing clustering; MATLAB’s built in fastTextWordEmbedding with 12 clusters). **(G)** Goodness-of-fit measured as the number of neurons with negative explained variance as a function of number of categories and across models.

**Figure S10:**
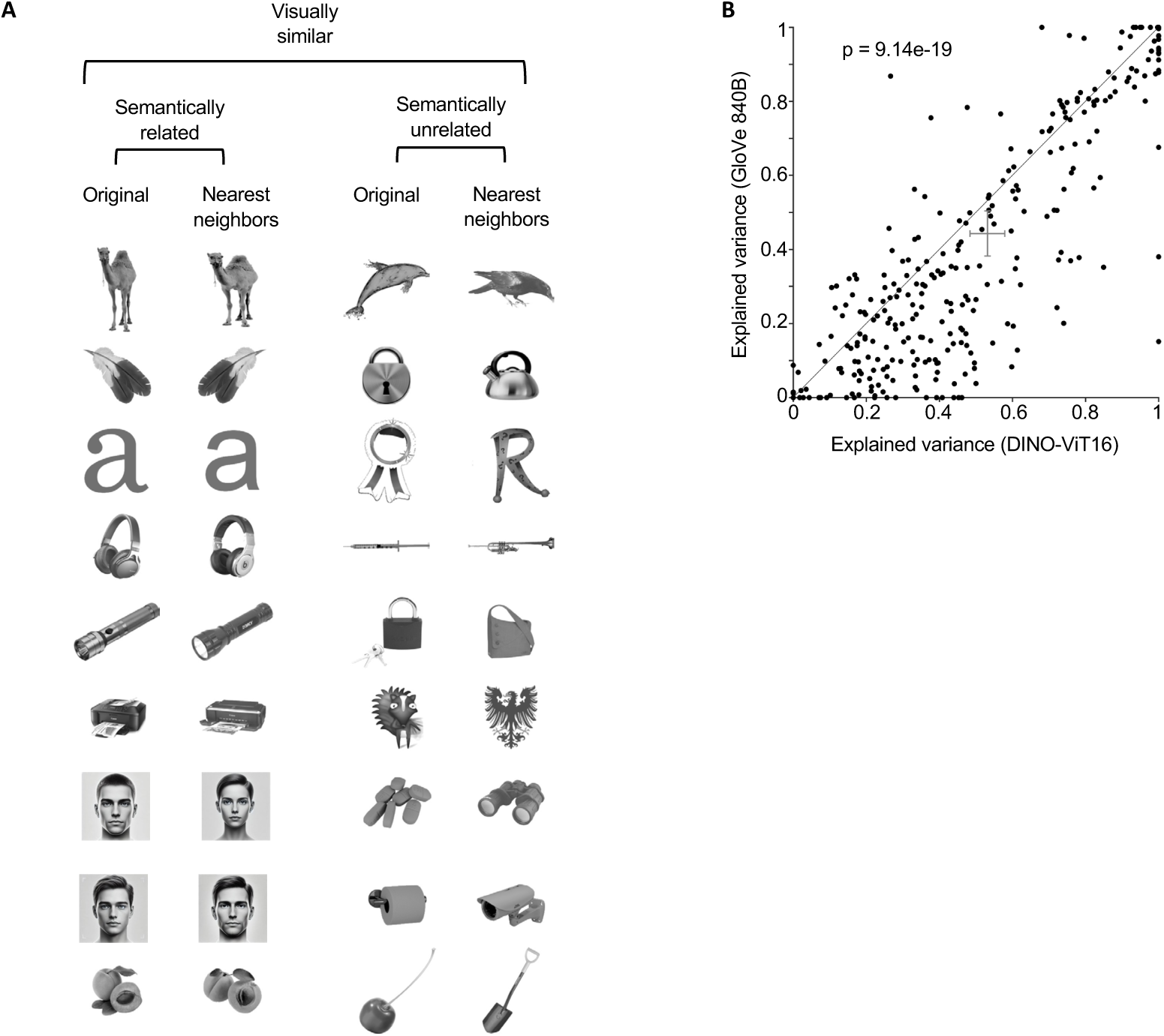
The response of axis-tuned VTC neurons is visual. [Note: The human faces in panel A of this figure have been replaced with synthetic faces generated by a diffusion model (*130*), in accordance with the bioarxiv policy on displaying human faces.] **(A)** Examination of the nearest neighbors of the 500 stimulus images in fc6 space. For 187/500 (∼40%) images the nearest neighbors were semantically unrelated despite being visually very similar. Shown are 18 **e**xamples of stimulus images (‘original’, columns one and three), 9 of which have semantically related nearest neighbors (column two) and 9 that have semantically unrelated nearest neighbors (column four). Regardless of semantic relatedness, all nearest neighbors are highly visually similar, indicating that fc6 is organized by visual features and not semantic associations. **(B)** We compared the neural variance explained between two types of embeddings: one that is purely visual by construction (DINO-ViT, a network trained in an unsupervised manner without labels (*72*)), and one that is purely semantic by construction (vector representation of the labels (*85*) – see supplementary methods). The visual model explained significantly more neural variance than the semantic model (Figure S10B; DINO-ViT = 53.12%, GloVe 840B = 44.23%, estimated out of sample, p = 9.14e-19 paired t-test).

## Supplementary tables

**Table S1.**
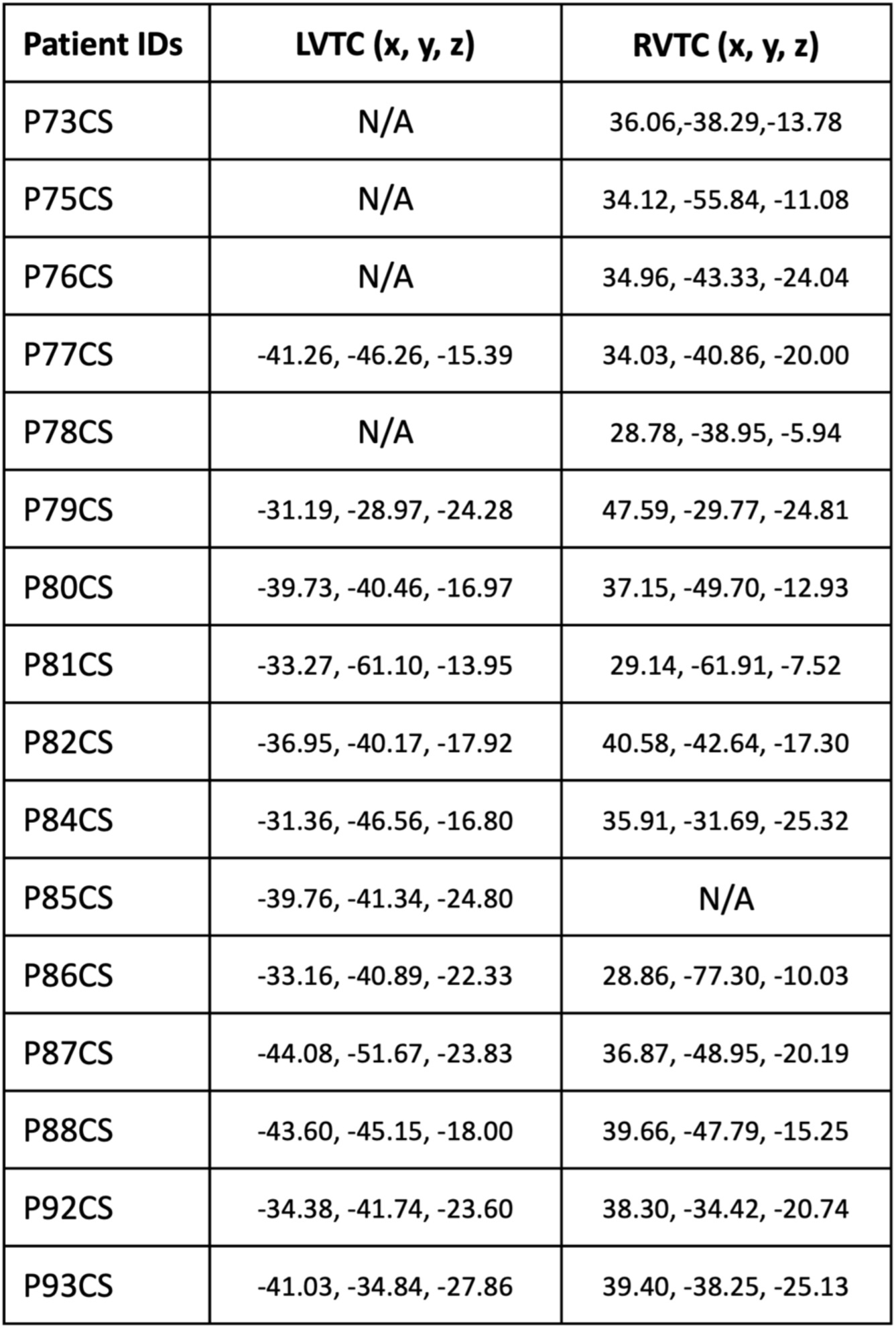
MNI coordinates of microwire bundles in which at least 1 VTC neuron was recorded. Related to Figure 1.

**Table S2.**
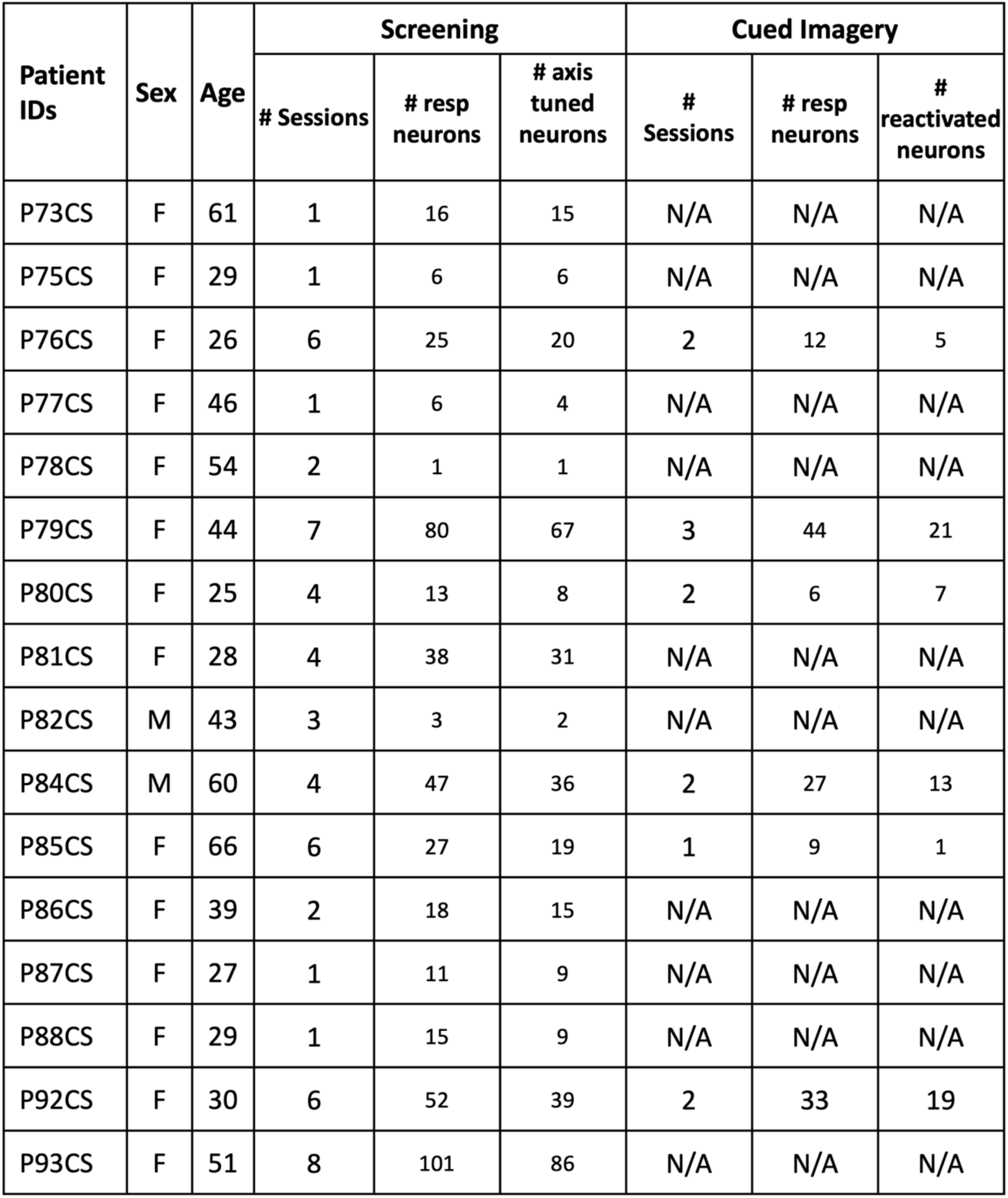
Number of sessions and neurons recorded. Summary of the number of neurons of each type recorded in each patient. In some subjects both Screening and Cued Imagery tasks were performed.

**Table S3.** Imagery correlations across different models. Pearson correlation between the projection value onto the axis computed during screening and firing rate during imagery for the preferred and orthogonal axes in all axis-tuned and reactivated neurons (n = 43), across all models. Firing rate during imagery correlates significantly with projection value on to preferred axis but not with projection onto the orthogonal axis regardless of the embedding used.

| Model | Corr. pref | P<sub>pref</sub> | Corr. ortho | P<sub>ortho</sub> |
| --- | --- | --- | --- | --- |
| AlexNet | 0.203 | 0 | -0.084 | 0.94 |
| VGGface | 0.087 | 0.052 | -0.135 | 0.994 |
| VGG16 | 0.203 | 0 | 0.027 | 0.321 |
| VGG19 | 0.190 | 0 | 0.014 | 0.423 |
| Cornet-Z | 0.234 | 0 | 0.005 | 0.471 |
| Cornet-S | 0.218 | 0 | 0.063 | 0.129 |
| Cornet-R/RT | 0.209 | 0 | -0.019 | 0.641 |
| Eigen Model | 0.129 | 0.008 | 0.0067 | 0.423 |
| DiNO-ViT16 | 0.135 | 0.01 | 0.014 | 0.374 |

**Table S4.** Imagery correlations across all layers of AlexNet. Pearson correlation between the projection value onto the axis computed during screening and firing rate during imagery for the preferred and orthogonal axes in all axis-tuned and reactivated neurons (n = 43), across all layers of AlexNet. Fc6 shows the highest correlation of projection value and firing rate during imagery.

| AlexNet layer | Corr. pref | P<sub>pref</sub> | Corr. ortho | P<sub>ortho</sub> |
| --- | --- | --- | --- | --- |
| 'conv1' | 0.067 | 0.103 | -0.154 | 0.998 |
| 'relu1' | 0.095 | 0.031 | 0.088 | 0.06 |
| 'norm1' | 0.126 | 0.013 | -0.098 | 0.97 |
| 'pool1' | 0.138 | 0.004 | 0.042 | 0.206 |
| 'conv2' | 0.158 | 0.002 | -0.109 | 0.978 |
| 'relu2' | 0.1856 | 0.001 | 0.069 | 0.086 |
| 'norm2' | 0.171 | 0 | -0.050 | 0.826 |
| 'pool2' | 0.127 | 0.009 | -0.132 | 0.994 |
| 'conv3' | 0.169 | 0.001 | 0.072 | 0.103 |
| 'relu3' | 0.200 | 0 | 0.073 | 0.096 |
| 'conv4' | 0.179 | 0 | 0.130 | 0.006 |
| 'relu4' | 0.202 | 0 | 0.036 | 0.248 |
| 'conv5' | 0.166 | 0 | 0.062 | 0.14 |
| 'relu5' | 0.172 | 0 | 0.040 | 0.247 |
| 'pool5' | 0.159 | 0.002 | -0.076 | 0.907 |
| 'fc6' | 0.203 | 0 | 0.051 | 0.189 |
| 'relu6' | 0.191 | 0 | 0.076 | 0.087 |
| 'fc7' | 0.151 | 0.003 | 0.053 | 0.157 |
| 'relu7' | 0.145 | 0.005 | -0.067 | 0.883 |
| 'fc8' | 0.136 | 0.005 | -0.050 | 0.807 |
| 'prob' | 0.039 | 0.246 | -0.016 | 0.62 |

